# Transcriptome Analysis Identified *SPP1+* Monocytes as a Key in Extracellular Matrix Formation in Thrombi

**DOI:** 10.1101/2024.05.28.594130

**Authors:** Takaya Kitano, Tsutomu Sasaki, Takahiro Matsui, Masaharu Kohara, Kotaro Ogawa, Todo Kenichi, Hajime Nakamura, Yuri Sugiura, Yuki Shimada, Shuhei Okazaki, Junichi Iida, Kohki Shimazu, Eiichi Morii, Manabu Sakaguchi, Masami Nishio, Masaru Yokoe, Haruhiko Kishima, Hideki Mochizuki

**Author notes:** Corresponding: Tsutomu Sasaki Department of Neurology, Osaka University Graduate School of Medicine, Osaka, Japan.

## Abstract

Thrombi follow various natural courses. They are known to become harder over time and may persist long-term; some of them can also undergo early spontaneous dissolution and disappearance. Hindering thrombus stability may contribute to the treatment of thrombosis and the prevention of embolisms. However, the detailed mechanisms underlying thrombus maturation remain unknown. Using RNA sequencing, we revealed the transcriptional landscape of thrombi retrieved from the cerebral vessels and identified *SPP1* as a hub gene related to extracellular matrix formation. Immunohistochemistry confirmed the expression of osteopontin in monocytes/macrophages in the thrombi, particularly in older thrombi. Single-cell RNA sequencing of thrombi from the pulmonary artery revealed increased communication between *SPP1*-high monocytes/macrophages and fibroblasts. These data suggest that *SPP1*-high monocytes/macrophages play a crucial role in extracellular matrix formation in thrombi and provide a basis for new antithrombotic therapies targeting thrombus maturation.

**Teaser:** *SPP1+* monocytes play a key role in thrombus maturation, which can be a potential target for novel antithrombotic therapies.

## Introduction

Thromboembolic diseases account for 1 in 4 deaths worldwide (*1*). These diseases can be classified into two groups, based on whether the thrombus is formed under high- or low-pressure systems (*2*), with intracardiac (cardiac atria) and venous thrombi accounting for most of the latter. Cardiogenic embolism, which can be caused by the embolism of an intracardiac thrombus, is the most severe type of ischemic stroke. Prevention of cardiogenic embolism is of great social interest because the number of patients with atrial fibrillation, the largest risk factor for cardiogenic embolism, is rapidly increasing (*3*). Venous thrombi primarily form in the lower extremities and can be symptomatic. In some patients, they can embolize the pulmonary artery, which can be fatal. Anticoagulants are primarily used to prevent and treat thromboembolic diseases under low-pressure systems. Anticoagulants can decrease the risk of cardioembolic stroke in patients with atrial fibrillation; however, an approximately 40% stroke risk remains even upon treatment with anticoagulants (*4*). In addition, patients may experience bleeding complications (*5*). Therefore, a more effective antithrombotic therapy with a low risk of bleeding is desirable.

Thrombi detected in patients can either disappear or persist. More than half (63–89%) of thrombi found in atrial appendages disappear without symptomatic embolic events, while some patients develop cerebral embolism (*6-9*). Moreover, 35% of surgery-associated venous thrombi resolve spontaneously within 72 hours, while some extend to involve the proximal veins (*10, 11*). Furthermore, half of pulmonary embolisms resolve within a few weeks, while thrombi persist; chronic thromboembolic pulmonary hypertension (CTEPH) occurs in approximately 5% of patients after pulmonary embolism (*11, 12*). Recently, dynamic changes occurring in thrombi over time have been noted (*13, 14*), which are presumably involved in determining the natural course of thrombi. As time passes after a deep venous thrombus is formed, the proportion of red blood cells (RBCs) decreases, and the level of extracellular matrix (ECM) proteins, such as collagen, increases (*14*). Thrombi can become stiffer and more resistant to thrombolysis over time (*15, 16*). We have previously reported that older thrombi are more resistant to reperfusion therapy in patients with cerebral embolism (*17*). Hence, it can be assumed that there are conflicting processes within a formed thrombus: some to make it stiff and stable and others to dissolve it. However, the mechanisms underlying these processes are not completely understood.

Here, we report the transcriptional differences between thrombi retrieved from cerebral vessels and peripheral blood. Our data revealed several major processes occurring in the thrombus, including ECM formation and inflammatory and anti-inflammatory responses. We identified *SPP1* as an upregulated hub gene for ECM formation in the thrombus. Furthermore, using single-cell RNA sequencing data, our results suggest that *SPP1*-high monocytes/macrophages (MCs/MPs) are key players in ECM formation in thrombi. Collectively, our data provide a basis for developing new antithrombotic therapies that modify the natural history of thrombi.

## Results

### RNA expression in thrombi differs from that in blood

Mechanical thrombectomy is an endovascular surgery procedure that has rapidly developed over the last decade to recanalize occluded cerebral vessels in patients with acute ischemic stroke. In this study, we compared the RNA expression profile in three thrombi retrieved from cerebral vessels via mechanical thrombectomy with that in simultaneously sampled blood (Fig. 1A). All patients were diagnosed with a cardiogenic embolism. The heatmap in Fig. 1B shows a substantial difference in RNA expression between the thrombus and blood. Unbiased clustering detected sample differences (Fig. 1C). Compared to that in the blood, a total of 1,121 genes were significantly upregulated and 693 were downregulated in the thrombus (Fig. 1D). The top 20 upregulated differentially expressed genes (DEGs) included pro-inflammatory chemokines such as *CXCL8* and *CCL2*, consistent with a previous report (Fig. 1E) (*18*). We also found that genes related to the ECM, such as *FN1* and *SPP1*, were upregulated in the thrombi.

**Fig. 1.**
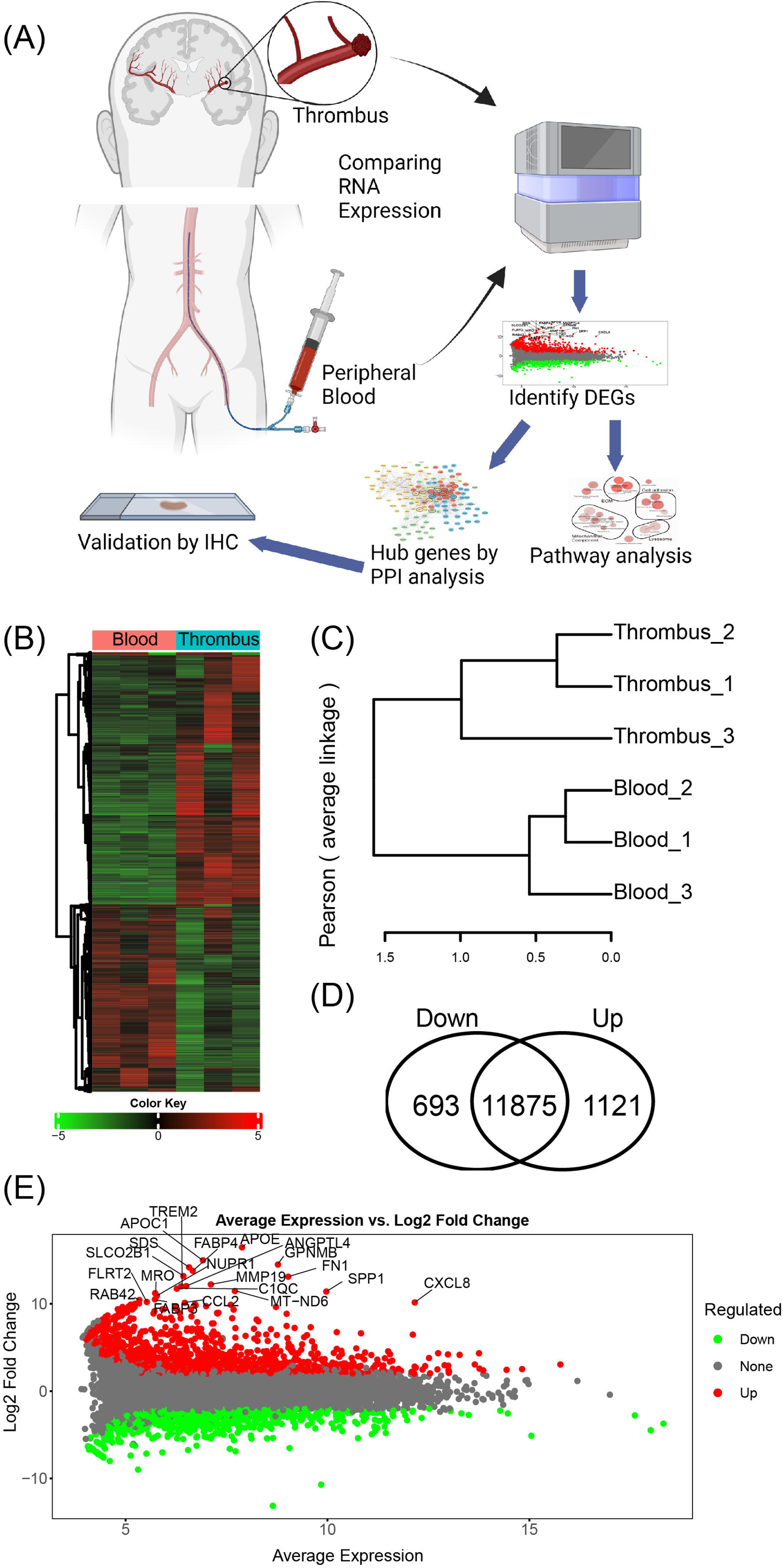
Comparison of gene expression profiles between the thrombi and blood. (A) Summary of the analysis results. DEGs, differentially expressed genes; IHC, immunohistochemistry; PPI, protein-protein interaction. (B) Heatmaps of the gene expression patterns. (C) Unsupervised hierarchical clustering of the samples. (D) Number of upregulated and downregulated genes in the thrombi compared to that in the blood. (E) MA plot of the genes. Genes upregulated in thrombi are shown in red, and genes with the top 20 highest fold-changes are annotated.

### Gene sets related to ECM were enriched in thrombi

The results of the gene set enrichment analysis are shown in Fig. 2A as network plots. Biological process analysis revealed clusters, such as tissue morphogenesis and ECM organization. Cellular component analysis revealed clusters of ECM, cell adhesion, and mitochondrial components. Molecular function analysis revealed clusters such as chemokine activity, cell adhesion, and ECM constituents. In all analyses, pathways related to the ECM were significantly enriched. Genes related to ECM were highly expressed in thrombi (Supplementary Fig. S1)

**Fig. 2.**
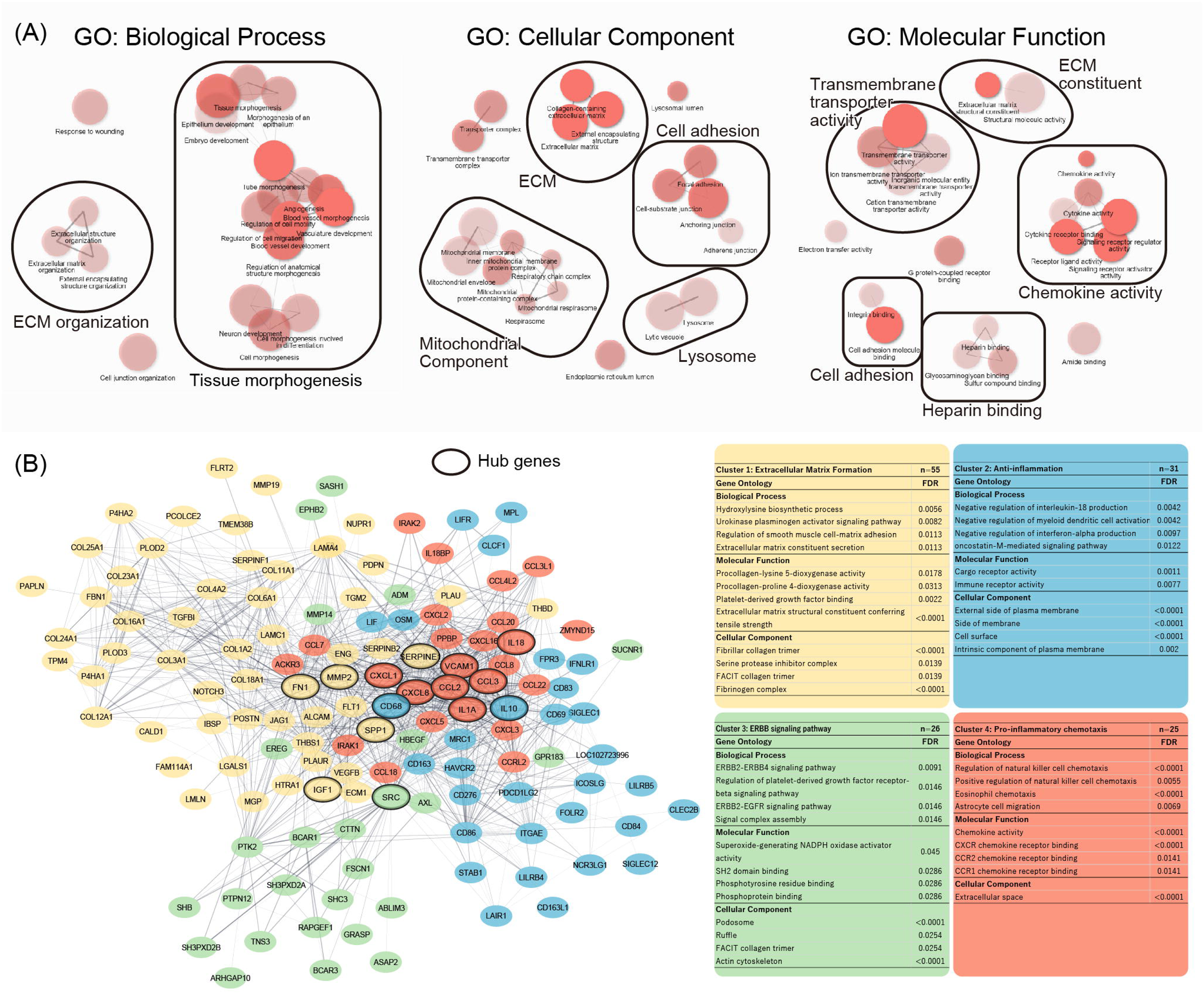
Gene set enrichment and protein-protein interaction analysis. (A) Gene set enrichment analysis results. The top 20 upregulated pathways in each category are visualized. Pathways sharing genes are connected by edges. The width of the edges represents the number of shared genes. The pathways are manually clustered and annotated. (B) Protein-protein interaction network. Genes belonging to the top 4 clusters out of the 963 upregulated genes identified in the STRING database are visualized. Clustering was performed using MCL and each cluster is represented by different colors. The thickness of each edge represents the level of confidence. Hub genes are identified through their Matthews Correlation Coefficients and are highlighted in bold circles. Gene set enrichment analysis results according to each cluster are shown as a table on the right and are arranged according to strength. Cluster annotation was performed manually. FDR, false discovery rate.

### Protein-protein interaction (PPI) analysis identified SPP1 as one of the hub genes

Among the 1,121 upregulated DEGs, 1,038 genes were identified in the STRING database after excluding the mitochondrial genes. We then identified 23 hub genes using CytoHubba (Supplementary Fig. S2). Unbiased MCL clustering identified 278 clusters using 963 genes. The top four clusters, which included the most genes among the identified clusters, are visualized in Fig. 2B. Cluster 1 included 55 genes associated with ECM formation. Cluster 2 included 31 genes related to the anti-inflammatory response. Cluster 3 included 26 genes related to the ERBB signaling pathway. Cluster 4 included 25 genes related to pro-inflammatory cytokines. Fifteen hub genes were included in the top four clusters.

We identified five hub genes (*FN1*, *MMP2*, *IGF1*, *SERPINE1*, and *SPP1*) in the ECM formation cluster, as this cluster included the most genes and pathways related to ECM were highly enriched in the gene set enrichment analysis (Fig. 2A). *FN1* encodes for fibronectin, a glycoprotein involved in cell adhesion, migration, wound healing, and embryonic development. Fibronectin is vital for bleeding control (*19*). *MMP2* encodes matrix metalloproteinase-2, which is involved in the degradation and remodeling of various ECM components. It was reported that *MMP2* inactivation prevented thrombosis and prolonged bleeding time (*20*). IGF-1 (insulin-like growth factor 1) is essential in the insulin signaling pathway and is a key growth factor involved in various processes such as cell proliferation, maturation, differentiation, and survival (*21*). It also regulates fibroblast growth and extracellular matrix deposition (*22*). *SERPINE1* encodes plasminogen activator inhibitor-1; its deficiency causes abnormal bleeding (*23*). *SPP1* encodes osteopontin (OPN), a secreted multifunctional glycophosphoprotein that plays an important role in physiological and pathophysiological processes (*24*). OPN drives immune responses under ischemic conditions and induces neutrophil and macrophage infiltration (*25-27*). It was recently 1, a potential target for α9β1, a preventing arterial thrombosis (*28-30*). Therefore, we decided to investigate whether OPN is present in thrombi retrieved from cerebral vessels and to determine how OPN is associated with the clinical characteristics of patients who underwent mechanical thrombectomy.

### Osteopontin is expressed by monocytes/macrophages in thrombi

We first validated the expression of *SPP1* via RT-qPCR using five pairs of samples, adding two other pairs to the samples used for RNA sequencing. In the thrombi retrieved from the cerebral artery, the expression of *SPP1* and its known receptor (*CD44*) was elevated compared to that in the blood. Additionally, *TIMP1* which is modulated by OPN and related to ECM (*31*), was also elevated.

We then performed immunohistochemical staining of paraffin-embedded thrombus samples using three anti-OPN antibodies. All antibodies validated the presence of OPN in thrombi (Supplementary Fig. S3). All the observed samples were positive for OPN, but the extent varied (Fig. 3B, 3C). In one thrombus, some regions showed strong positivity, whereas others did not (Fig. 3D). We also observed OPN+ cell aggregation (where OPN+ cells were observed as a cluster) when the thrombus was strongly positive for OPN. Thus, we classified the thrombi into OPN-high or OPN-low according to the presence or absence of OPN+ cell aggregation. Double staining revealed that OPN was not expressed by neutrophils that are defined as cells with a lobulated nucleus and positive for neutrophil elastase (Fig. 3E). OPN was mainly expressed by MC/MPs in the thrombi (Fig. 3F).

**Fig. 3.**
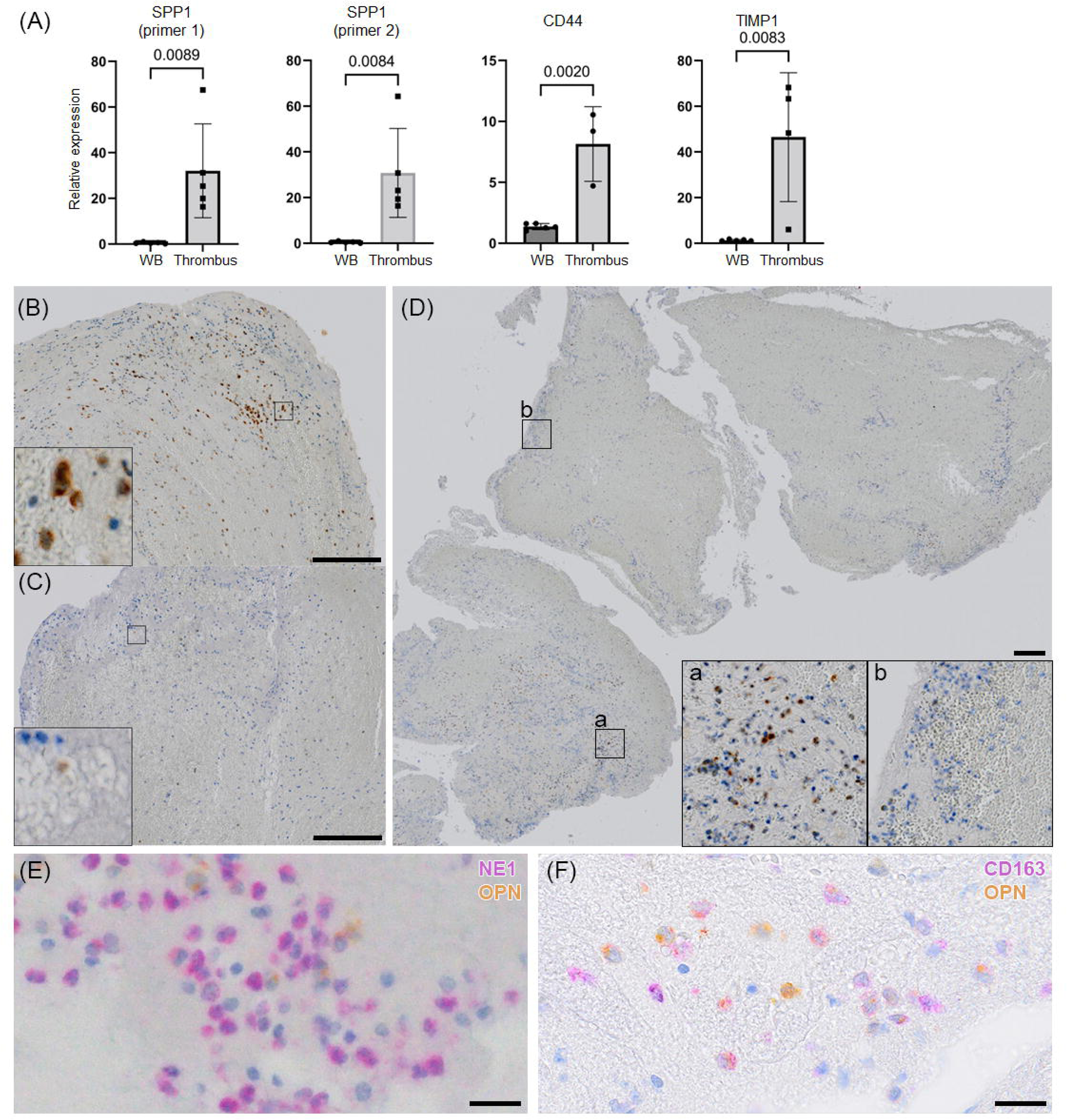
Immunohistochemical staining shows the presence of OPN in thrombi. (A) Gene expression as measured using qPCR. (B¬¬–D) Immunohistochemical staining for OPN using rabbit polyclonal antibodies against the N-terminal part of human OPN (18625). Boxed areas are magnified in the panel. Bar = 200 μ image of a thrombus with strong positivity for OPN. The aggregation of OPN+ cells was observed. C, Thrombus with weak positivity for OPN. D, OPN+ cells unevenly distributed in one thrombus. OPN, osteopontin. (E) Double-staining against neutrophil elastase 1 (purple) and OPN (yellow). Bar = 20 μm (F) Double-staining against CD163 (purple) and OPN (yellow).

### Osteopontin expression is more observed in older thrombi than in fresh thrombi

Next, we analyzed 66 thrombi retrieved during mechanical thrombectomy for cerebral embolisms (Supplementary Fig. S4). Among the 66 samples, 40 were OPN-high and 26 were OPN-low. We then compared the features of the thrombi based on OPN expression. Fresh thrombi were less common among OPN-high samples (46% vs. 3%, P < 0.001; Fig. 4A). No significant differences were observed in the proportions of red RBCs, fibrin, or platelets. The density of MCs/MPs was higher in the OPN-high samples (Fig. 4B). As the density of MC/MPs is higher in older thrombi than in fresh ones (*17, 32*), these observations support the idea that OPN-high thrombi tend to be older than OPN-low thrombi.

**Fig. 4.**
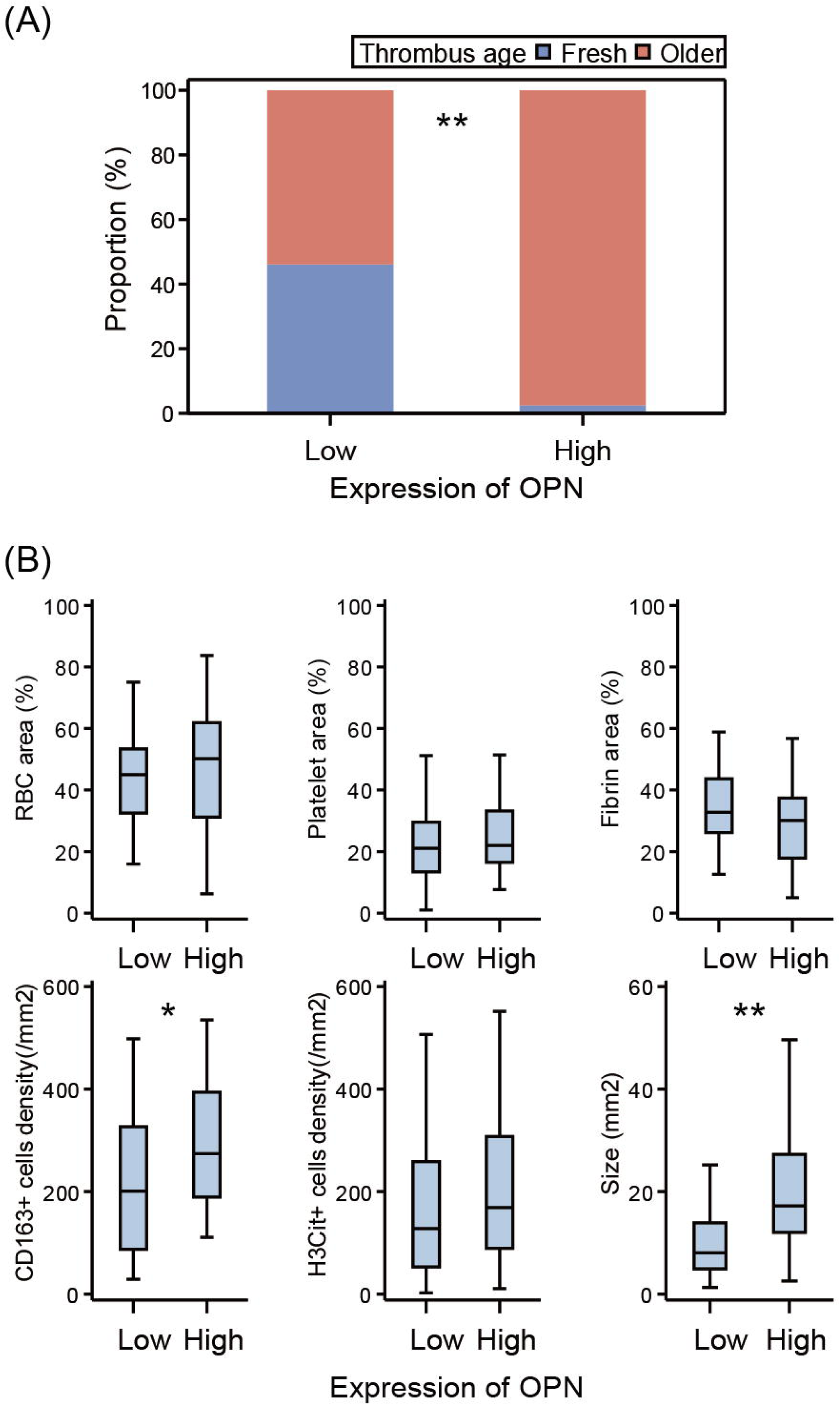
Comparison of thrombus features according to OPN expression. (A) Fresh thrombi are more common among the OPN-low thrombi (P < 0.001) than older thrombi. (B) Comparison of thrombus components, CD163+ cell density, and the degree of NETosis. *P < 0.05, **P < 0.01.

### Osteopontin expression and clinical characteristics

To determine the association between OPN expression in thrombi and the clinical characteristics of patients, we compared patient backgrounds according to OPN expression (Table 1). The proportion of patients with atrial fibrillation was marginally higher (54% vs. 78%, P = 0.060), the level of brain natriuretic peptide was marginally higher (82.5 vs. 225 pg/mL, P = 0.054), and the cardiothoracic ratio was significantly higher (57% vs. 62%, P = 0.023) in patients with OPN-high thrombi. The proportion of stroke subtypes also differed significantly across OPN expression (P = 0.043, Fig. 5A); the proportion of cardioembolic stroke was higher among patients with OPN-high thrombi. We then investigated the effect of OPN expression on reperfusion quality. There was no significant difference in the time to reperfusion after arterial puncture across OPN expression levels (P = 0.26, Fig. 5B), and no significant difference was observed in the number of passes before successful reperfusion (P = 0.17, Fig. 5C).

**Fig 5.**
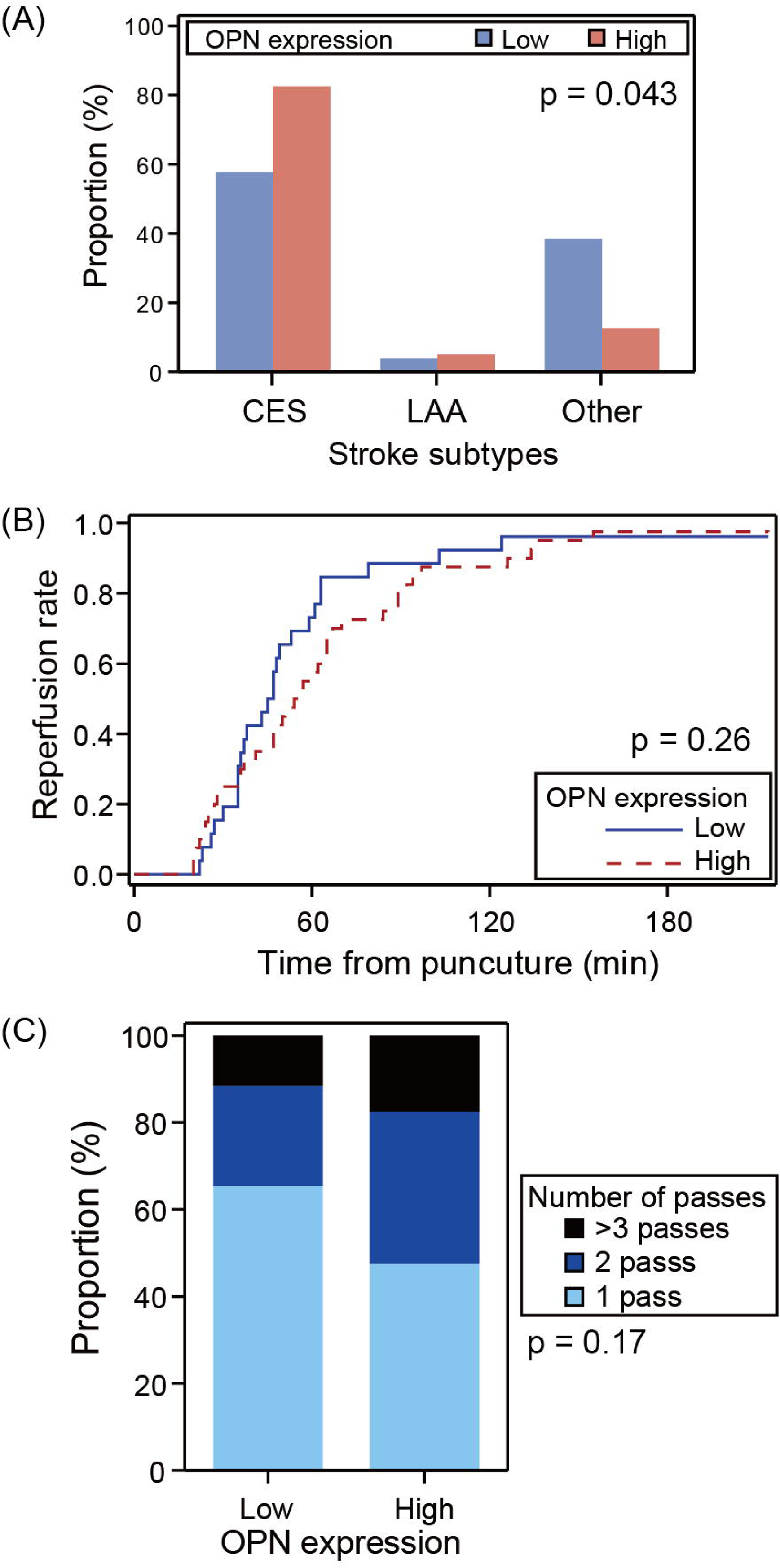
Associations between clinical characteristics and OPN expression in the thrombus. (A) Proportion of patients according to stroke subtype. (B) Cumulative rate of successful reperfusion (expanded treatment in cerebral ischemia ≥ 2b) after puncture. (B) Proportion of the number of passes before successful reperfusion.

**Table 1.** Background characteristics according to OPN expression in the thrombus.

|  | OPN low<br>(n = 26) | OPN high<br>(n = 40) | P | miss |
| --- | --- | --- | --- | --- |
| Age, yrs | 78 (71–83) | 79.5 (66–85) | 0.80 | 0 |
| Male | 13 (50%) | 19 (48%) | 1 | 0 |
| Hypertension | 16 (62%) | 23 (58%) | 0.80 | 0 |
| Diabetes | 5 (19%) | 5 (13%) | 0.50 | 0 |
| Dyslipidemia | 10 (39%) | 14 (35%) | 0.80 | 0 |
| Atrial fibrillation | 14 (54%) | 31 (78%) | 0.060 | 0 |
| Antiplatelets | 8 (31%) | 11 (28%) | 0.79 | 0 |
| Anticoagulants | 7 (27%) | 12 (30%) | 1 | 0 |
| Modified Rankin Scale | 0 (0–2) | 0.5 (0–2) | 0.30 | 0 |
| NIHSS score | 16.5 (12–24) | 17.5 (13–22.5) | 0.77 | 0 |
| Leukocyte count, /μL | 8,110 (6,350–11,500) | 7,130 (5,280–9,110) | 0.097 | 0 |
| Platelet count, $10^4/\mu$L | 21.5 (19.2–23.8) | 18.4 (15.7–22.2) | 0.021 | 0 |
| C-reactive protein, mg/dL | 0.16 (0.05–0.40) | 0.265 (0.11–1.66) | 0.28 | 0 |
| Fibrinogen, mg/dL | 285 (253–374) | 321.5 (289–398) | 0.35 | 27 |
| D-dimer, μg/mL | 2.48 (1–4.61) | 1.7 (0.8–3.18) | 0.194 | 4 |
| Brain Natriuretic Peptide, pg/mL | 82.5 (43–253) | 225 (94–555.7) | 0.054 | 21 |
| Cardio-thoracic ratio, % | 57 (52–63) | 62 (58–65) | 0.023 | 2 |
| ASPECTS | 9.5 (7.5–10) | 9 (7–10) | 0.83 | 5 |
| Tandem lesion | 1 (4%) | 2 (5%) | 1 | 0 |
| Hyperdense MCA sign | 13 (57%) | 20 (59%) | 1 | 9 |
| Susceptibility vessel sign | 6 (60%) | 6 (75%) | 0.64 | 48 |
| rt-PA administration | 10 (39%) | 15 (38%) | 1 | 0 |
| Onset-to-puncture time, min | 239 (130–383) | 187 (105–362) | 0.49 | 0 |

Abbreviations: ASPECTS, Alberta Stroke Program Early CT Score; MCA, middle cerebral artery; NIHSS, National Institute of Health Stroke Scale; OPN, osteopontin; rtPA, recombinant tissue-plasminogen activator

### SPP1-high monocytes/macrophages in thrombi are related to ECM formation

After confirming that *SPP1* was expressed by MC/MPs in stroke thrombi, we investigated whether *SPP1* was also expressed in thrombi retrieved from patients with other thromboembolic diseases and the role of *SPP1+* MC/MPs in thrombi. We used single-cell RNA sequencing data of thrombi from patients with CTEPH (*33*) to characterize MCs/MPs that express *SPP1* in the thrombus. After unbiased clustering (Supplementary Fig. S5), we successfully identified MCs/MPs (Fig. 6A, 6B). *SPP1* was mainly expressed by MC/MPs in these samples. Four subclusters of MCs/MPs were identified (Fig. 6C), and *SPP1* was listed as one of the top ten highly expressed genes in subcluster 2 (Fig. 6D). The expression of hub genes identified via PPI analysis is shown in Fig. 6E. *CD68* was highly expressed in subcluster 2, whereas *CXCL8* and *CCL3* were highly expressed in subcluster 1. Gene set enrichment analysis revealed that pathways related to the extracellular matrix are upregulated in subcluster 2 (Fig. 6E, red circle). Pathways related to lysosomes (Fig. 6E, pink and light blue circles) were also upregulated in subcluster 2. In addition, pathways related to inflammation, such as the cellular response to tumor necrosis factor, response to interleukin-1, and chemotaxis, were enriched in subcluster 1 (Supplementary Fig. S6A).

**Fig. 6.**
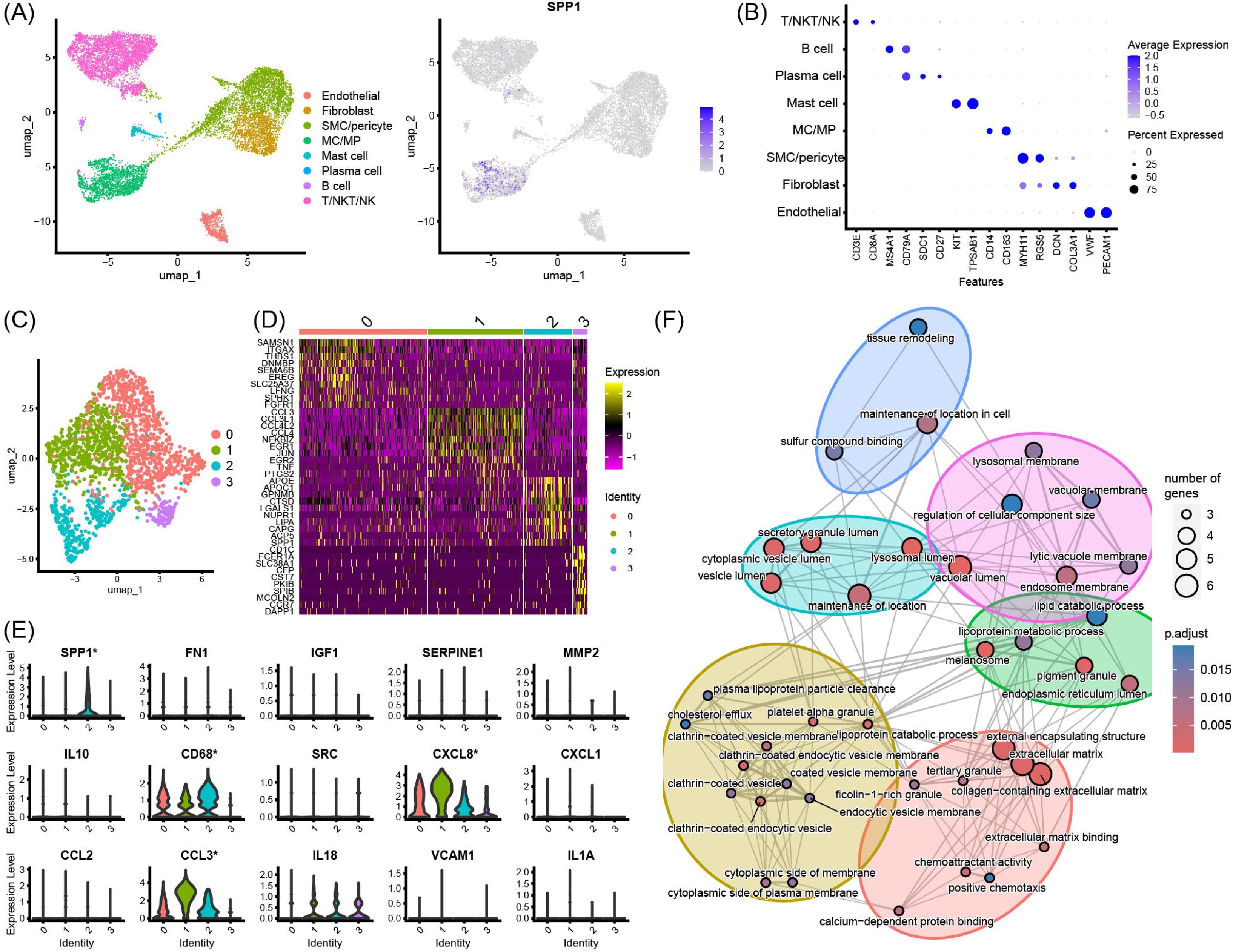
Analysis of single-cell RNA sequencing data of thrombi from patients with chronic thromboembolic pulmonary hypertension. (A) UMAP plot visualizing the cell type and feature plot of the expression of SPP1. (B) Expression of specific marker genes. (C) Subclusterng of the MC/MPs. Four subclusters are identified. (D) Heatmap of the top 10 differentially expressed genes according to each subcluster of MC/MPs. SPP1 is highly expressed in subcluster 2. (E) Expression of hub genes identified by the PPI in the thrombus. *P_val_adj < 0.0001. (F) Gene set enrichment analysis results. The pathways upregulated in subcluster 2 MC/MPs are visualized. Pathways sharing genes are connected by edges, and the width of the edges represents the number of shared genes. Clustering was performed automatically.

To understand the role of subcluster 2 (*SPP1*-high MC/MPs) in thrombus formation, we performed a ligand-receptor interaction analysis using CellChat. Dense communication between MCs/MPs and fibroblasts, which are the major source of ECM, was inferred in the thrombi from patients with CTEPH (Fig. 7A). Among the immune cells found in the thrombus, MCs/MPs in subcluster 2 had the highest number and strength of inferred ligand-receptor interactions as the sender between fibroblasts (Fig. 7B). When MCs/MPs were assigned to the sender, the ligand-receptor pairs *SPP1*-*ITGAV*_*ITGB1*, *SPP1*-*ITGA8*_*ITGB1*, and *SPP1*-*ITGAV*_*ITGB5* were inferred to be the main communicators between subcluster 2 MCs/MPs and fibroblasts (Fig. 7C, 7D). In addition, ligand-receptor pairs of *PDGFB*-*PDGFRB* were inferred between subcluster 1 MC/MPs and fibroblasts (Supplementary Fig. S6B).

**Fig. 7.**
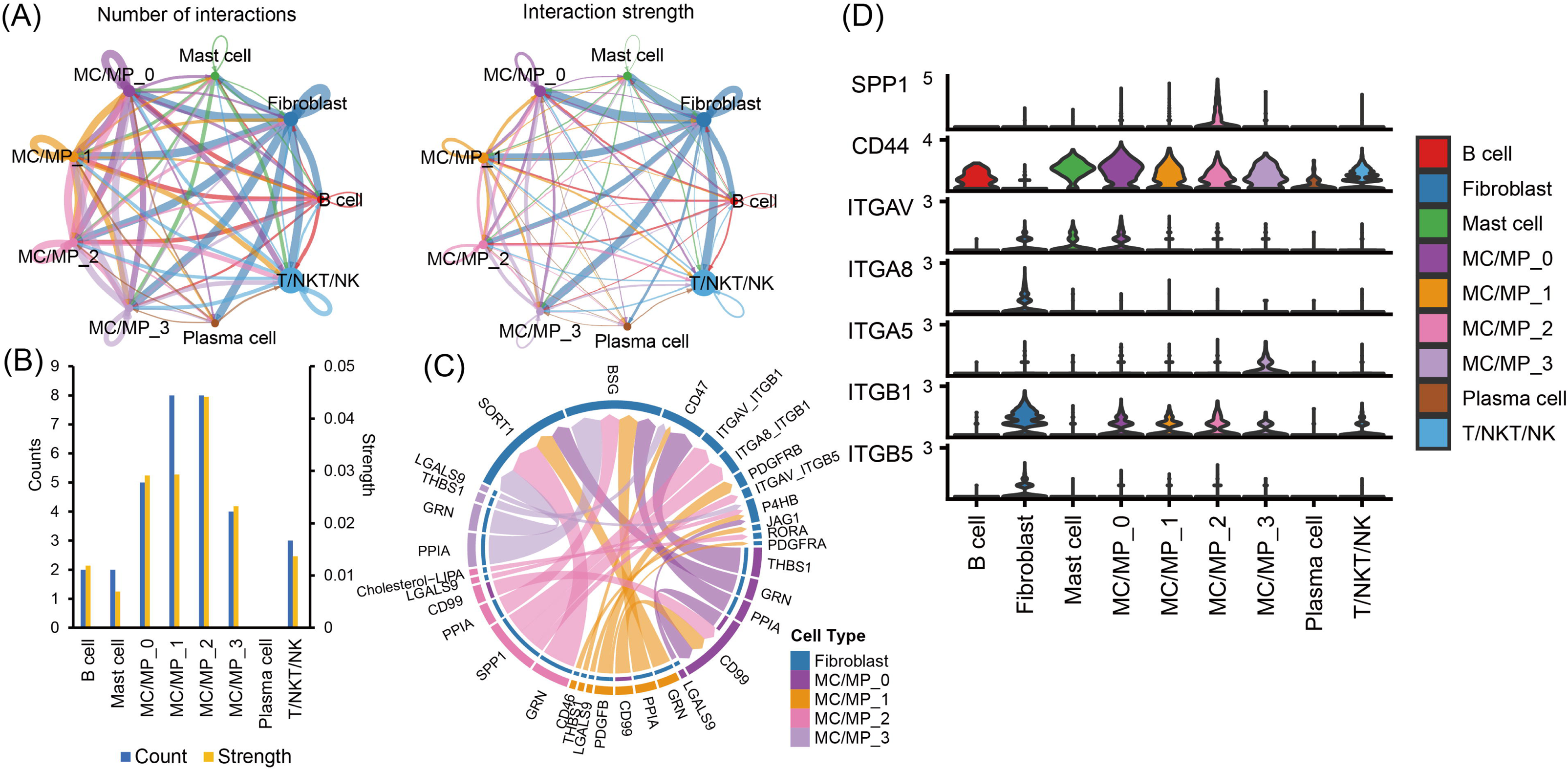
Ligand-receptor interaction analysis using single-cell RNA sequencing data from thrombi of patients with chronic thromboembolic pulmonary hypertension. (A) Network plots of the inferred numbers and strengths of ligand-receptor interactions. (B) Inferred numbers and strengths of ligand-receptor interactions with fibroblasts according to cluster. Fibroblasts were set as the receiver. (C) Chord diagram to visualize all the inferred ligand-receptor pairs from the MC/MPs subclusters to fibroblasts. (D) Expression of genes in the SPP1 signaling pathway.

### SPP1-high monocytes/macrophages are expanded in murine venous vessel walls after thrombosis induction

As MC/MPs are presumed to migrate into the thrombus through the vessel wall (*34*), we hypothesized that *SPP1*-high MC/MPs may be present in the vessel wall after thrombosis. We investigated the expression of *SPP1* by MC/MPs in the venous vessel walls of mice with and without deep vein thrombosis (DVT) (Supplementary Fig. S7A– S7D). Our results showed that the subcluster of MC/MPs with high *SPP1* expression was expanded in the DVT group (Supplementary Fig. S7E).

## Discussion

In this study, we compared RNA expression in thrombi from acute cerebral infarctions with that in simultaneously collected peripheral blood. In addition to pro-inflammatory responses, which have been reported previously (*18*), our findings revealed an anti-inflammatory response and ECM formation in thrombi. *SPP1* is one of the hub genes upregulated in thrombi, and the expression of its translation product, OPN, was observed particularly in older thrombi retrieved from the cerebral vessels. Furthermore, single-cell RNA sequencing data from thrombi in patients with CTEPH revealed that *SPP1*-high MC/MPs likely contributed to ECM formation. These results elucidate the transcriptional landscape that occurs after thrombus formation and could contribute to the development of new antithrombotic drugs that target thrombus maturation.

*SPP1+* macrophages have recently been identified as key cell types promoting tissue fibrosis (*35, 36*). In line with these reports, the results of gene set enrichment analysis suggest that subcluster 2 MCs/MPs (*SPP1*-high) are related to the ECM in the thrombi of patients with CTEPH. This was supported by the results of our ligand-receptor interaction analysis. Integrin αV, an OPN receptor expressed by fibroblasts, is a key molecule in fibrotic diseases (*37, 38*). The interaction between OPN and integrins V 1 and V 5 has been reported to contribute to the adhesion of smooth muscle and fibroadipogenic progenitor cells (*36, 39*); OPN is also chemotactic for smooth muscle cells (*40*). Hence, *SPP1-*high MC/MPs may promote ECM formation by recruiting fibroblasts to the thrombus. Nonetheless, our results do not exclude the possibility that other subclusters of MCs/MPs are involved in ECM formation in the thrombus. Subcluster 1 MCs/MPs have been shown to promote fibrogenesis through *PDGFB*-*PDGFRB* interactions. In addition, the *THBS1*-*CD47* interaction, which has been reported to promote fibrosis (*41*), was also found in other subclusters. Further investigations are required to clarify the role of heterogeneous MC/MPs in ECM formation in thrombi.

OPN+ cells were not evenly distributed throughout the thrombus in thrombi retrieved from cerebral vessels. They were often densely aggregated in certain areas, particularly around the periphery of the thrombus. These aggregations of OPN+ cells are presumed to occur sometime after thrombus formation, as they are less common in fresh thrombi. In DVT, macrophages are presumed to increase migrating from the vascular wall to the thrombus during venous thrombosis (*17, 34, 42*). The present study showed that *SPP1*-high MC/MPs were expanded in the venous wall after thrombosis induction. Moreover, a recent study revealed that *SPP1*+ macrophages are expanded in the left atrial tissue of patients with persistent atrial fibrillation and could be targets for immunotherapy in atrial fibrillation (*43*). Therefore, some of the OPN+ MCs/MPs observed in thrombi from patients with cardioembolic stroke may have migrated from the left atrial wall. The observation in the present study that patients without atrial fibrillation had fewer OPN-high thrombi supports this hypothesis. Hence, the migration of OPN+ MCs/MPs may be a potential target for new antithrombotic therapies.

Intracardiac thrombi are frequently encountered in clinical practice, and the prevalence of left atrial appendage thrombus formation is estimated to be between 5% and 27% in patients who have not previously received anticoagulant therapy (*44*). Patients with intracardiac thrombi are at an elevated risk of developing cardiogenic embolisms. Thus, convenient and cost-effective screening methods for intracardiac thrombi are required. Recently, OPN has been recognized as a biomarker for vascular diseases (*24*). The plasma concentration of OPN was found to be higher in patients with atrial fibrillation than in those without and is correlated with the extent of atrial fibrosis (*45*). Using unbiased proteomics, Mühlen et al. reported that OPN levels in the urine could be a biomarker for venous thrombosis and pulmonary embolism (*46*). It was also reported that plasma OPN levels are higher among patients with venous thrombosis than in those without (*47*). OPN is cleaved by thrombin into two halves. In this study, we showed that cleaved OPN (N-half) was present in a thrombus retrieved from a cerebral vessel. Considering that the presence of cleaved OPN implies a thrombogenic state, the level of OPN, especially cleaved OPN, in the peripheral blood may have potential value as a biomarker to predict intracardiac thrombus and future embolic events in patients with atrial fibrillation.

In the present study, thrombi retrieved from the cerebral vessels were found to contain neutrophils, consistent with previous studies (*32, 48, 49*). However, neutrophils could not be identified in the single-cell RNA sequencing data of thrombi harvested from patients with CTEPH. Experimental findings in a mouse inferior vena cava ligation model indicated that neutrophils are abundant in the initial stages of thrombus formation; however, their proportion decreased over time, with the proportion of MC/MPs increasing later on (*50*). Therefore, neutrophils may disappear from the CTEPH thrombus with age.

This study has several limitations. First, some RNA integrity numbers (RIN) were not sufficiently high in the thrombus samples. RINs in thrombus samples are commonly low, and acellular RINs tend to have lower RNA quantity and quality (*51*). We chose samples with a relatively high RIN for RNA sequencing and validated the results using qPCR after adding other samples; however, careful interpretation of the results is needed. Second, the sample size of the clinical data may not have been sufficiently large to detect differences in reperfusion quality.

In conclusion, transcriptional responses, including ECM formation, inflammatory response, and anti-inflammatory response, were identified in thrombi in this study. Our results collectively suggest that *SPP1+* MC/MPs play a key role in ECM formation in thrombi and may be a potential target for new antithrombotic therapies that modify thrombus maturation.

## Materials and Methods

### Sample collection and total RNA isolation

This study was approved by the Institutional Review Board of the Toyonaka Municipal Hospital, and all participants provided written informed consent. The clinical backgrounds of the participants are shown in Supplemental Table S1. Thrombi retrieved from five patients with acute ischemic stroke were stored in RNAlater (Thermo Fisher, Waltham, MA, USA) overnight at 4°C immediately after mechanical thrombectomy. The thrombi were then transferred to a -80°C freezer until subsequent analyses. Whole blood was sampled from the patients via the femoral arterial sheath during the mechanical thrombectomy and stored using a PAXgene RNA blood collection tube (762165, BD Biosciences, San Jose, CA, USA) in a -80°C freezer until the subsequent steps. Total RNA was extracted from each thrombus using a ReliaPrep RNA tissue MiniPrep System (Z6110, Promega, Madison, WI, USA) and from whole blood using a PAXgene Blood RNA Kit (762164, Qiagen, Hilden, Germany) according to the respective manufacturer’s instructions.

### Library preparation and sequencing

The purity and quantity of the isolated total RNA were assessed using a NanoDrop 2000 spectrophotometer (Thermo Fisher). The RIN was assessed using Bioanalyzer 2100 (Agilent, Santa Clara, CA, USA). The three thrombus RNA samples with the highest RINs and their paired blood RNA samples were subjected to subsequent analyses. The RINs for the thrombus samples were 8.0, 6.8, and 4.5, and those for the paired blood samples were 7.2, 7.1, and 7.7. The total RNA samples were subjected to library preparation using an Illumina TruSeq stranded Total RNA Library Prep Kit with Ribo-Zero Globin (Illumina, San Diego, CA, USA) according to the manufacturer’s protocol. RNA libraries were subjected to 100-bp paired-end sequencing on a NovaSeq 6000 system (Illumina) with a median of 40 million reads.

### RNA sequencing data processing, mapping, and counting

Sequence quality was assessed using FastQC (v0.12.1, Babraham Bioinformatics), and reads were trimmed using Trimmomatic (v. 0.36) (*52*). The reads were mapped to GRCh38 using HISAT2 software (ver. 2.2.1) (*53*). Mapped reads were counted using the feature Counts (v2.0.3) (*54*). All these procedures were performed on the Galaxy platform (*55*) and default parameters were used.

### Differentially expressed genes and gene set enrichment analysis

DEGs between the thrombi and blood were identified using an integrated browser application, iDEP1.1 (*56*). Minimal counts of 1.5/million in at least two libraries were set in the preprocessing data interface, and the count data were subjected to a variance-stabilizing transform for clustering. A heat map was generated to include the top 2,000 genes. Unsupervised hierarchical clustering was performed using 1-Pearson correlation and average linkage. DEGs were identified using the iDEP built-in DESeq2 package (*57*) with a threshold of false discovery rate < 0.01 and a minimum absolute value of fold-change > 4.

Gene set enrichment analysis was performed using the DEGs defined above to determine the most enriched gene ontology (GO) pathways in terms of biological processes, cellular components, and molecular functions. Genes with a false discovery rate > 0.6 were removed from the pathway analysis. The top 20 pathways were displayed as a network. Nodes were connected if they shared 20% or more genes. All parameters not specified above were left as the default values. Pathway clusters were manually annotated.

### Protein-protein interaction (PPI) analysis

The PPIs between all DEGs upregulated in the thrombus, except for mitochondrial genes, were analyzed using the STRING database (v12.0) (*58*). A minimal interaction score of 0.4 was set as the default. Hub genes were identified using Cytoscape (v3.10.1) software and the CytoHubba plugin (v0.1) (*59*). DEGs with the highest Matthews Correlation Coefficient scores were considered hub genes. The DEGs were clustered using MCL with a default inflation parameter of 3; GO enrichment analysis was performed on these clusters. Clusters were manually annotated based on the enriched pathways and visualized using Cytoscape software.

### Quantitative real-time PCR analysis

We performed RT-qPCR to amplify and detect *SPP1* and its receptor *CD44* from five pairs of thrombus and blood RNA samples, comprising three pairs used for RNA-seq analysis and two additional pairs. First, cDNA was prepared by reverse-transcribing 400 pg of total RNA using Superscript III and random primers (Thermo Fisher Scientific). The resulting cDNA was used for real-time PCR analysis using a Platinum SYBR Green qPCR SuperMix (Thermo Fisher Scientific). Next, 100 reverse transcription products and standard plasmids were subjected to real-time PCR analysis (QuantStudio 7 Flex Real-Time PCR System; Applied Biosystems) using human β-actin as an internal control. The following PCR program was used: 10 min of denaturation at 95°C, and then 40 cycles of 95°C for 15 s, 58°C for 30 s, and 72°C for 30 s. The primers used are listed in Supplementary Table S2.

### Subjects for the histological analysis of the thrombus

We examined 168 consecutive patients who underwent mechanical thrombectomy for acute ischemic stroke between January 2015 and December 2019 at two tertiary referral hospitals with comprehensive stroke centers in Japan (Osaka University Hospital, Osaka; Osaka General Medical Center, Osaka). Thrombus specimens were available for 76 patients. Patients with left ventricular assist devices, atherosclerotic intracranial stenosis, or cerebral artery dissection were excluded. Finally, 66 patients with thrombi retrieved during mechanical thrombectomy for cerebral embolisms were included in this study (Supplementary Fig. S4). Clinical data were also collected from the included patients. The detailed methods for clinical data collection were as previously described (*17*).

### Immunohistochemical staining and histological analysis

Thrombus samples were fixed in 10% neutral-buffered formalin and embedded in paraffin. To identify the presence of osteopontin (OPN), a product of *SPP1,* three primary antibodies were used: mouse monoclonal antibodies (10011, IBL, Gunma, Japan), rabbit polyclonal antibodies (25715-1-AP, Proteintech, Rosemont, IL, USA), and rabbit polyclonal antibodies against the N-terminal region of human OPN (18625, IBL). Immunohistochemical staining was performed using a Roche Ventana BenchMark GX autostainer (Ventana Medical Systems, Tucson, AZ, USA), according to the manufacturer’s instructions. We stained human gallbladder and kidney samples as positive controls to determine the optimal antibody concentrations. The stained slides were captured as digital images using a VS200 Slide Scanner (Olympus, Tokyo, Japan). The age, size, and components of the thrombi and the extent of NETosis were evaluated as previously described (*17*). Thrombus age was evaluated based on hematoxylin and eosin (H&E) staining and positivity for anti-alpha-smooth muscle actin (*60*). Thrombus size and RBC proportion were measured using H&E staining, fibrin was detected using phosphotungstic acid-hematoxylin staining, and platelets were subjected to immunohistochemical staining for CD42b. The density of MC/MPs was determined via staining for CD163, and the extent of NETosis was determined using H3Cit staining. A thrombus was considered OPN-high when the aggregation of OPN+ cells was observed. In addition, four samples were subjected to double-staining with anti-OPN and anti-neutrophil elastase antibodies or with anti-OPN and anti-CD163 antibodies.

### Single-cell RNA sequencing data analysis of CTEPH thrombi

We obtained publicly available single-cell RNA sequencing data of thrombi collected from patients with CTEPH, which were deposited by Rajagopal et al. (PRJNA929967) (*33*). Filtered matrices were loaded into the R package Seurat (v.5.0) (*61*), and cells with less than 200 features, more than 9,000 features, less than 300 UMIs, more than 60,000 UMIs, or more than 15% mitochondrial gene fractions were removed. Normalization was performed using the R package sctransform (*62*), followed by the integration workflow of Seurat. Principal component analysis was performed on the integrated data, and the top 30 principal components were used to cluster the cells. The FindNeighbors and FindClusters functions in Seurat were used to identify cell clusters with a resolution of 1.0. The FindAllMarkers function in Seurat was used to identify the markers for each cluster. Clusters were manually annotated using known lineage markers and several clusters were combined as needed. MC/MPs were defined as cells with a high expression of *CD14* or *CD163*. MC/MPs were subjected to subclustering using a resolution of 0.2. DEGs from each subcluster were calculated using the FindAllMarkers function, and the top 10 upregulated genes were used for the DoHeatmap function. Pathway enrichment analysis was performed using the enrichGO function in the R package clusterProfiler (*63*). We set the adjusting P method to FDR, the threshold to 0.05, the minimum gene set size to 10, and the maximum gene set size to 600. Pathways that included fewer than three genes were excluded. The top 40 pathways with the smallest P values were visualized using the Emapplot function. The bound pathways were automatically clustered using the Emapplot function.

The receptor-ligand interaction between each subcluster of MCs/MPs and fibroblasts was analyzed using CellChat (v. 2.1) (*64*). Normalized data were used for each condition, and cell types with fewer than 10 cells were excluded. Interaction strength refers to the probability of communication between a given ligand and receptor. It was calculated as the degree of cooperativity/interactions derived from the law of mass action and the degree to which the ligands and receptors are expressed.

### Single-cell RNA sequencing data analysis of cells from murine venous walls

We obtained single-cell RNA sequencing data from cells of the vein wall of a mouse DVT model, which were deposited by Zhou et al. (PRJNA916965) (*65*). Cells with less than 800 features, more than 8,000 features, less than 1000 UMIs, more than 40,000 UMIs, or more than 20% mitochondrial gene fractions were removed from the analysis. Data analysis was performed as above, except that the top 15 principal components were used and that clustering was performed with a resolution of 0.2. Identified MCs/MPs were subjected to subclustering using the top 15 principal components and a resolution of 0.6.

### Statistical analysis of clinical data

Continuous variables were reported as the median and interquartile range (IQR), while categorical variables were reported as numbers and percentages. Continuous variables were compared using the Wilcoxon rank-sum test. Categorical variables were compared using Fisher’s exact test. The cumulative rate of successful reperfusion was evaluated using the Kaplan–Meier method. Statistical significance was set P < 0.05. All analyses were performed using SAS Studio software (SAS 9.4; SAS Institute Inc., Cary, NC, USA).

## Author contributions

Conceptualization: TK, TS, TM Methodology: TK, TS

Investigation: TK, TS, MK, KO, KT, HN, YURS, YUKS, SO, JI, HS, MS, and MN

Visualization: TK, TS, TM, MK Supervision: TS, MY, HM

Writing—original draft: TK, TS

Writing—review & editing: EM, MY, HK, HM

## Data and materials availability

RNA sequencing data of the thrombi were deposited in the NIH Gene Expression Omnibus with accession ID PRJNA1099305. The clinical data are available upon request.

## References

1. A. M. Wendelboe, G. E. Raskob, Global Burden of Thrombosis. Circulation Research 118, 1340–1347 (2016).

2. K. T. Tan, G. Y. H. Lip, Red vs White Thrombi: Treating the Right Clot Is Crucial. Archives of internal medicine 163, 2534–2535 (2003).

3. H. Kamel, J. S. Healey, Cardioembolic Stroke. Circulation Research 120, 514–526 (2017).

4. R. G. Hart, L. A. Pearce, M. I. Aguilar, Meta-analysis: antithrombotic therapy to prevent stroke in patients who have nonvalvular atrial fibrillation. Annals of internal medicine 146, 857–867 (2007).

5. M. T. Kalathottukaren, C. A. Haynes, J. N. Kizhakkedathu, Approaches to prevent bleeding associated with anticoagulants: current status and recent developments. Drug Deliv Transl Res 8, 928–944 (2018).

6. L. J. Collins, D. I. Silverman, P. S. Douglas, W. J. Manning, Cardioversion of nonrheumatic atrial fibrillation. Reduced thromboembolic complications with 4 weeks of precardioversion anticoagulation are related to atrial thrombus resolution. Circulation 92, 160–163 (1995).

7. W. A. Jaber, D. L. Prior, M. Thamilarasan, R. A. Grimm, J. D. Thomas, A. L. Klein, C. R. Asher, Efficacy of anticoagulation in resolving left atrial and left atrial appendage thrombi: A transesophageal echocardiographic study. American heart journal 140, 150–156 (2000).

8. G. Corrado, G. Tadeo, S. Beretta, L. M. Tagliagambe, G. F. Manzillo, M. Spata, M. Santarone, Atrial thrombi resolution after prolonged anticoagulation in patients with atrial fibrillation. Chest 115, 140–143 (1999).

9. G. Y. Lip, C. Hammerstingl, F. Marin, R. Cappato, I. L. Meng, B. Kirsch, M. van Eickels, A. Cohen, Left atrial thrombus resolution in atrial fibrillation or flutter: Results of a prospective study with rivaroxaban (X-TRA) and a retrospective observational registry providing baseline data (CLOT-AF). American heart journal 178, 126–134 (2016).

10. V. V. Kakkar, C. T. Howe, C. Flanc, M. B. Clarke, NATURAL HISTORY OF POSTOPERATIVE DEEP-VEIN THROMBOSIS. The Lancet 294, 230–233 (1969).

11. C. Kearon, Natural History of Venous Thromboembolism. Circulation 107, I-22–I-30 (2003).

12. A. Ribeiro, P. Lindmarker, H. Johnsson, A. Juhlin-Dannfelt, L. Jorfeldt, Pulmonary embolism: one-year follow-up with echocardiography doppler and five-year survival analysis. Circulation 99, 1325–1330 (1999).

13. J. M. Nicklas, A. E. Gordon, P. K. Henke, Resolution of Deep Venous Thrombosis: Proposed Immune Paradigms. International journal of molecular sciences 21, (2020).

14. M. Nosaka, Y. Ishida, A. Kimura, T. Kondo, Time-dependent organic changes of intravenous thrombi in stasis-induced deep vein thrombosis model and its application to thrombus age determination. Forensic Sci Int 195, 143–147 (2010).

15. H. Xie, K. Kim, S. R. Aglyamov, S. Y. Emelianov, M. O’Donnell, W. F. Weitzel, S. K. Wrobleski, D. D. Myers, T. W. Wakefield, J. M. Rubin, Correspondence of ultrasound elasticity imaging to direct mechanical measurement in aging DVT in rats. Ultrasound Med Biol 31, 1351–1359 (2005).

16. K. P. Mercado-Shekhar, R. T. Kleven, H. Aponte Rivera, R. Lewis, K. B. Karani, H. J. Vos, T. A. Abruzzo, K. J. Haworth, C. K. Holland, Effect of Clot Stiffness on Recombinant Tissue Plasminogen Activator Lytic Susceptibility in Vitro. Ultrasound Med Biol 44, 2710–2727 (2018).

17. T. Kitano, Y. Hori, S. Okazaki, Y. Shimada, T. Iwamoto, H. Kanki, S. Sugiyama, T. Sasaki, H. Nakamura, N. Oyama, T. Hoshi, G. Beck, H. Takai, S. Matsubara, H. Mizuno, H. Nishimura, R. Tamaki, J. Iida, J. Iba, M. Uno, H. Kishima, H. Fushimi, S. Hattori, S. Murayama, E. Morii, M. Sakaguchi, Y. Yagita, T. Shimazu, H. Mochizuki, K. Todo, An Older Thrombus Delays Reperfusion after Mechanical Thrombectomy for Ischemic Stroke. Thromb Haemost, (2021).

18. R. A. Campbell, A. Vieira-de-Abreu, J. W. Rowley, Z. G. Franks, B. K. Manne, M. T. Rondina, L. W. Kraiss, J. J. Majersik, G. A. Zimmerman, A. S. Weyrich, Clots Are Potent Triggers of Inflammatory Cell Gene Expression: Indications for Timely Fibrinolysis. Arteriosclerosis, thrombosis, and vascular biology 37, 1819–1827 (2017).

19. Y. Wang, A. Reheman, C. M. Spring, J. Kalantari, A. H. Marshall, A. S. Wolberg, P. L. Gross, J. I. Weitz, M. L. Rand, D. F. Mosher, J. Freedman, H. Ni, Plasma fibronectin supports hemostasis and regulates thrombosis. J Clin Invest 124, 4281–4293 (2014).

20. S. Momi, E. Falcinelli, S. Giannini, L. Ruggeri, L. Cecchetti, T. Corazzi, C. Libert, P. Gresele, Loss of matrix metalloproteinase 2 in platelets reduces arterial thrombosis in vivo. The Journal of experimental medicine 206, 2365–2379 (2009).

21. M. Obradovic, S. Zafirovic, S. Soskic, J. Stanimirovic, A. Trpkovic, D. Jevremovic, E. R. Isenovic, Effects of IGF-1 on the Cardiovascular System. Curr Pharm Des 25, 3715–3725 (2019).

22. Y. Yin, Y. Han, C. Shi, Z. Xia, IGF-1 regulates the growth of fibroblasts and extracellular matrix deposition in pelvic organ prolapse. Open Med (Wars) 15, 833–840 (2020).

23. W. P. Fay, A. C. Parker, L. R. Condrey, A. D. Shapiro, Human plasminogen activator inhibitor-1 (PAI-1) deficiency: characterization of a large kindred with a null mutation in the PAI-1 gene. Blood 90, 204–208 (1997).

24. Z. S. Y. Lok, A. N. Lyle, Osteopontin in Vascular Disease. Arteriosclerosis, thrombosis, and vascular biology 39, 613–622 (2019).

25. G. S. Lee, H. F. Salazar, G. Joseph, Z. S. Y. Lok, C. M. Caroti, D. Weiss, W. R. Taylor, A. N. Lyle, Osteopontin isoforms differentially promote arteriogenesis in response to ischemia via macrophage accumulation and survival. Laboratory Investigation 99, 331–345 (2019).

26. C. L. Duvall, D. Weiss, S. T. Robinson, F. M. F. Alameddine, R. E. Guldberg, W. R. Taylor, The Role of Osteopontin in Recovery from Hind Limb Ischemia. Arteriosclerosis, thrombosis, and vascular biology 28, 290–295 (2008).

27. A. Koh, A. P. da Silva, A. K. Bansal, M. Bansal, C. Sun, H. Lee, M. Glogauer, J. Sodek, R. Zohar, Role of osteopontin in neutrophil function. Immunology 122, 466–475 (2007).

28. N. Dhanesha, M. K. Nayak, P. Doddapattar, M. Jain, G. D. Flora, S. Kon, A. K. Chauhan, 9 1 inhibits arterial thrombosis in mice. Blood 135, 857–861 (2020).

29. A. Brill, Integrin α β 9 1: a new target to fight thrombosis. Blood 135, 787–788 (2020).

30. N. Nishimichi, F. Higashikawa, H. H. Kinoh, Y. Tateishi, H. Matsuda, Y. Yokosaki, Polymeric osteopontin employs integrin alpha9beta1 as a receptor and attracts neutrophils by presenting a de novo binding site. J Biol Chem 284, 14769–14776 (2009).

31. T. Sabo-Attwood, M. E. Ramos-Nino, M. Eugenia-Ariza, M. B. Macpherson, K. J. Butnor, P. C. Vacek, S. P. McGee, J. C. Clark, C. Steele, B. T. Mossman, Osteopontin modulates inflammation, mucin production, and gene expression signatures after inhalation of asbestos in a murine model of fibrosis. The American journal of pathology 178, 1975–1985 (2011).

32. E. Laridan, F. Denorme, L. Desender, O. Francois, T. Andersson, H. Deckmyn, K. Vanhoorelbeke, S. De Meyer, Neutrophil extracellular traps in ischemic stroke thrombi: NETs in Stroke. Annals of Neurology 82, (2017).

33. A. G. Viswanathan, H. F. Kirshner, N. Nazo, S. Ali, A. Ganapathi, I. Cumming, Y. Zhuang, I. Choi A., Warman, C. Jassal, S. Almeida-Peters, J. Haney, D. Corcoran, Y. R. Yu, S. Rajagopal, Single-Cell Analysis Reveals Distinct Immune and Smooth Muscle Cell Populations that Contribute to Chronic Thromboembolic Pulmonary Hypertension. Am J Respir Crit Care Med 207, 1358–1375 (2023).

34. C. L. McGuinness, J. Humphries, M. Waltham, K. G. Burnand, M. Collins, A. Smith, Recruitment of labelled monocytes by experimental venous thrombi. Thromb Haemost 85, 1018–1024 (2001).

35. K. Hoeft, G. J. L. Schaefer, H. Kim, D. Schumacher, T. Bleckwehl, Q. Long, B. M. Klinkhammer, F. Peisker, L. Koch, J. Nagai, M. Halder, S. Ziegler, E. Liehn, C. Kuppe, J. Kranz, S. Menzel, I. Costa, A. Wahida, P. Boor, R. K. Schneider, S. Hayat, R. Kramann, Platelet-instructed SPP1+ macrophages drive myofibroblast activation in fibrosis in a CXCL4-dependent manner. Cell Reports 42, 112131 (2023).

36. M. Fu, S. Shu, Z. Peng, X. Liu, X. Chen, Z. Zeng, Y. Yang, H. Cui, R. Zhao, X. Wang, L. Du, M. Wu, W. Feng, J. Song, Single-Cell RNA Sequencing of Coronary Perivascular Adipose Tissue From End-Stage Heart Failure Patients Identifies SPP1(+) Macrophage Subpopulation as a Target for Alleviating Fibrosis. Arteriosclerosis, thrombosis, and vascular biology 43, 2143–2164 (2023).

37. A. N. C. Henderson, T. D. Arnold, Y. Katamura, M. M. Giacomini, J. D. Rodriguez, J. H. McCarty A., Pellicoro, E. Raschperger, C. Betsholtz, P. G. Ruminski, D. W. Griggs, M. J. Prinsen, J. J. Maher, J. P. Iredale, A. Lacy-Hulbert, R. H. Adams, D. Sheppard, Targeting of α integrin identifies a core molecular pathway that regulates fibrosis in several organs. Nature Medicine 19, 1617–1624 (2013).

38. I. R. Murray, Z. N. Gonzalez, J. Baily, R. Dobie, R. J. Wallace, A. C. Mackinnon, J. R. Smith, S. N. Greenhalgh, A. I. Thompson, K. P. Conroy, D. W. Griggs, P. G. Ruminski, G. A. Gray, M. Singh, M. A. Campbell, T. J. Kendall, J. Dai, Y. Li, J. P. Iredale, H. Simpson, J. Huard, B. Péault, N. C. Henderson, αv integrins on mesenchymal cells regulate skeletal and cardiac muscle fibrosis. Nat Commun 8, 1118 (2017).

39. L. Liaw, M. P. Skinner, E. W. Raines, R. Ross, D. A. Cheresh, S. M. Schwartz, C. M. Giachelli, The adhesive and migratory effects of osteopontin are mediated via distinct cell surface integrins. Role of alpha v beta 3 in smooth muscle cell migration to osteopontin in vitro. J Clin Invest 95, 713–724 (1995).

40. L. Liaw, M. Almeida, C. E. Hart, S. M. Schwartz, C. M. Giachelli, Osteopontin promotes vascular cell adhesion and spreading and is chemotactic for smooth muscle cells in vitro. Circ Res 74, 214–224 (1994).

41. G. Wernig, S. Y. Chen, L. Cui, C. Van Neste, J. M. Tsai, N. Kambham, H. Vogel, Y. Natkunam, D. G. Gilliland, G. Nolan, I. L. Weissman, Unifying mechanism for different fibrotic diseases. Proc Natl Acad Sci U S A 114, 4757–4762 (2017).

42. E. Furukoji, T. Gi, A. Yamashita, S. Moriguchi-Goto, M. Kojima, C. Sugita, T. Sakae, Y. Sato, T. Hirai, Y. Asada, CD163 macrophage and erythrocyte contents in aspirated deep vein thrombus are associated with the time after onset: a pilot study. Thrombosis journal 14, 46 (2016).

43. M. Hulsmans, M. J. Schloss, I. H. Lee, A. Bapat, Y. Iwamoto, C. Vinegoni, A. Paccalet, M. Yamazoe, J. Grune, S. Pabel, N. Momin, H. Seung, N. Kumowski, F. E. Pulous, D. Keller, C. Bening, U. Green, J. K. Lennerz, R. N. Mitchell, A. Lewis, B. Casadei, O. Iborra-Egea, A. Bayes-Genis, S. Sossalla, C. S. Ong, R. N. Pierson, J. C. Aster, D. Rohde, G. R. Wojtkiewicz, R. Weissleder, F. K. Swirski, G. Tellides, G. Tolis, Jr., S. Melnitchouk, D. J. Milan, P. T. Ellinor, K. Naxerova, M. Nahrendorf, Recruited macrophages elicit atrial fibrillation. Science 381, 231–239 (2023).

44. M. Patel, X. Wei, K. Weigel, Z. M. Gertz, J. Kron, A. A. Robinson, C. R. Trankle, Diagnosis and Treatment of Intracardiac Thrombus. J Cardiovasc Pharmacol 78, 361–371 (2021).

45. R. Lin, S. Wu, D. Zhu, M. Qin, X. Liu, Osteopontin induces atrial fibrosis by activating Akt/GSK-3β/β-catenin pathway and suppressing autophagy. Life Sciences 245, 117328 (2020).

46. C. von Zur Mühlen, T. Koeck, E. Schiffer, C. Sackmann, P. Zürbig, I. Hilgendorf, J. Reinöhl, J. Rivera, A. Zirlik, C. Hehrlein, H. Mischak, C. Bode, K. Peter, Urine proteome analysis as a discovery tool in patients with deep vein thrombosis and pulmonary embolism. Proteomics Clin Appl 10, 574–584 (2016).

47. A. A. Memon, K. Sundquist, M. PirouziFard, J. L. Elf, K. Strandberg, P. J. Svensson, J. Sundquist, B. Zöller, Identification of novel diagnostic biomarkers for deep venous thrombosis. Br J Haematol 181, 378–385 (2018).

48. V. M. Tutino, S. Fricano, A. Chien, T. R. Patel, A. Monteiro, H. H. Rai, A. A. Dmytriw, L. D. Chaves, M. Waqas, E. I. Levy, K. E. Poppenberg, A. H. Siddiqui, Gene expression profiles of ischemic stroke clots retrieved by mechanical thrombectomy are associated with disease etiology. J Neurointerv Surg, (2022).

49. C. Ducroux, L. Di Meglio, S. Loyau, S. Delbosc, W. Boisseau, C. Deschildre, M. Ben Maacha, R. Blanc, H. Redjem, G. Ciccio, S. Smajda, R. Fahed, J. B. Michel, M. Piotin, L. Salomon, M. Mazighi, B. Ho-Tin-Noe, J. P. Desilles, Thrombus Neutrophil Extracellular Traps Content Impair tPA-Induced Thrombolysis in Acute Ischemic Stroke. Stroke 49, 754–757 (2018).

50. M. Nosaka, Y. Ishida, A. Kimura, T. Kondo, Time-dependent appearance of intrathrombus neutrophils and macrophages in a stasis-induced deep vein thrombosis model and its application to thrombus age determination. Int J Legal Med 123, 235–240 (2009).

51. V. M. Tutino, S. Fricano, K. Frauens, T. R. Patel, A. Monteiro, H. H. Rai, M. Waqas, L. Chaves, K. E. Poppenberg, A. H. Siddiqui, Isolation of RNA from Acute Ischemic Stroke Clots Retrieved by Mechanical Thrombectomy. Genes (Basel) 12, (2021).

52. A. M. Bolger, M. Lohse, B. Usadel, Trimmomatic: a flexible trimmer for Illumina sequence data. Bioinformatics 30, 2114–2120 (2014).

53. D. Kim, B. Langmead, S. L. Salzberg, HISAT: a fast spliced aligner with low memory requirements. Nature Methods 12, 357–360 (2015).

54. Y. Liao, G. K. Smyth, W. Shi, featureCounts: an efficient general purpose program for assigning sequence reads to genomic features. Bioinformatics 30, 923–930 (2013).

55. T. G. Community, The Galaxy platform for accessible, reproducible and collaborative biomedical analyses: 2022 update. Nucleic Acids Research 50, W345–W351 (2022).

56. S. X. Ge, E. W. Son, R. Yao, iDEP: an integrated web application for differential expression and pathway analysis of RNA-Seq data. BMC Bioinformatics 19, 534 (2018).

57. M. I. Love, W. Huber, S. Anders, Moderated estimation of fold change and dispersion for RNA-seq data with DESeq2. Genome Biology 15, 550 (2014).

58. D. Szklarczyk, A. L. Gable, D. Lyon, A. Junge, S. Wyder, J. Huerta-Cepas, M. Simonovic, N. T. Doncheva, J. H. Morris, P. Bork, L. J. Jensen, C. V. Mering, STRING v11: protein-protein association networks with increased coverage, supporting functional discovery in genome-wide experimental datasets. Nucleic Acids Res 47, D607–d613 (2019).

59. C. H. Chin, S. H. Chen, H. H. Wu, C. W. Ho, M. T. Ko, C. Y. Lin, cytoHubba: identifying hub objects and sub-networks from complex interactome. BMC Syst Biol 8 Suppl 4, S11 (2014).

60. S. Z. Rittersma, A. C. van der Wal, K. T. Koch, J. J. Piek, J. P. Henriques, K. J. Mulder, J. P. Ploegmakers, M. Meesterman, R. J. de Winter, Plaque instability frequently occurs days or weeks before occlusive coronary thrombosis: a pathological thrombectomy study in primary percutaneous coronary intervention. Circulation 111, 1160–1165 (2005).

61. Y. Hao, T. Stuart, M. H. Kowalski, S. Choudhary, P. Hoffman, A. Hartman, A. Srivastava, G. Molla, S. Madad, C. Fernandez-Granda, R. Satija, Dictionary learning for integrative, multimodal and scalable single-cell analysis. Nature Biotechnology, (2023).

62. C. Hafemeister, R. Satija, Normalization and variance stabilization of single-cell RNA-seq data using regularized negative binomial regression. Genome Biology 20, 296 (2019).

63. G. Yu, L. G. Wang, Y. Han, Q. Y. He, clusterProfiler: an R package for comparing biological themes among gene clusters. Omics 16, 284–287 (2012).

64. S. Jin, M. V. Plikus, Q. Nie, CellChat for systematic analysis of cell-cell communication from single-cell and spatially resolved transcriptomics. bioRxiv, 2023.2011.2005.565674 (2023).

65. E. DeRoo, T. Zhou, H. Yang, A. Stranz, P. Henke, B. Liu, A vein wall cell atlas of murine venous thrombosis determined by single-cell RNA sequencing. Communications Biology 6, 130 (2023).

