## Supplementary material for "Transcriptome Analysis Identified *SPP1+* Monocytes as a Key in Extracellular Matrix Formation in Thrombi"

Takaya Kitano *et al.*

**This PDF file includes:**

Figs. S1 to S7

Tables S1 to S2

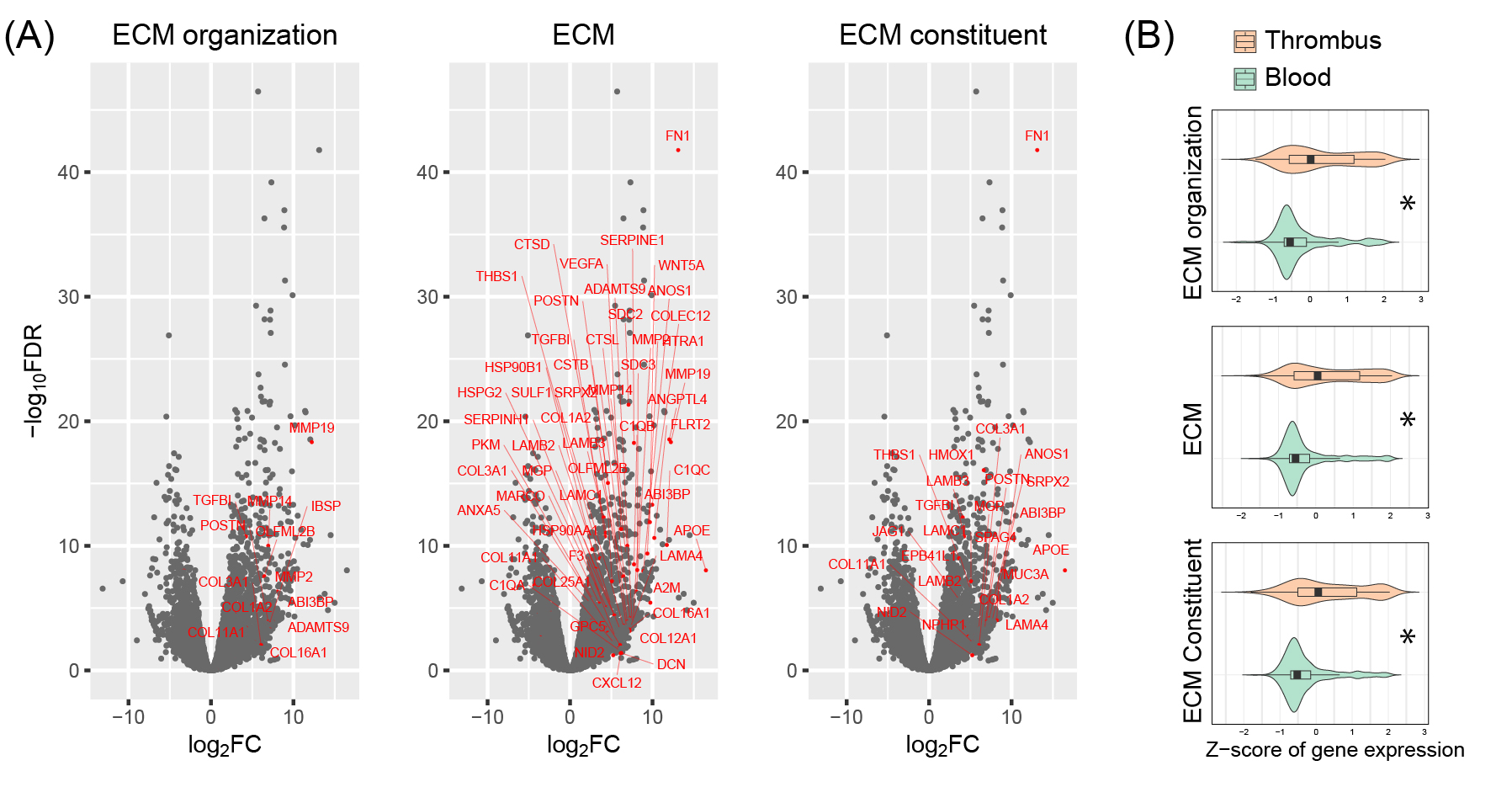

**Fig. S1. Genes related to ECM are highly expressed in thrombi. (A)** Volcano plots for gene expression in thrombi compared with blood. Gene sets related to ECM were generated based on the clusters found in the GO analysis (ECM organization, ECM, and ECM constituent). The genes included in each gene set are highlighted in red, and genes with log_2_FC>5 and/or -log_10_FDR>5 are labeled. **(B)** Z-score of the normalized gene expression included in each gene sets are compared according to the samples. * for p<0.001.

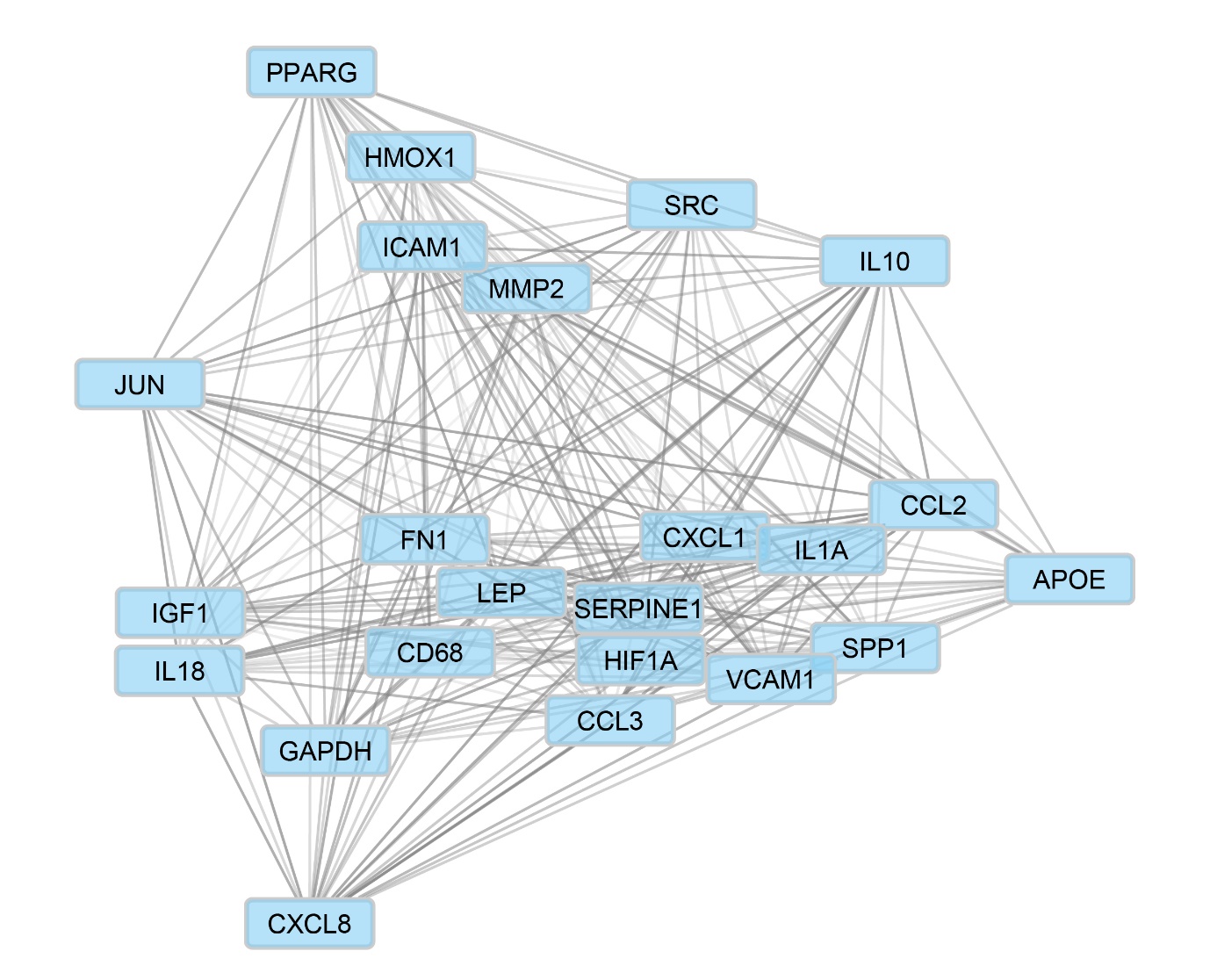

**Fig. S2. Hub genes identified using protein-protein interaction analysis.** The thicknesses of the edges represent the strength of the interaction.

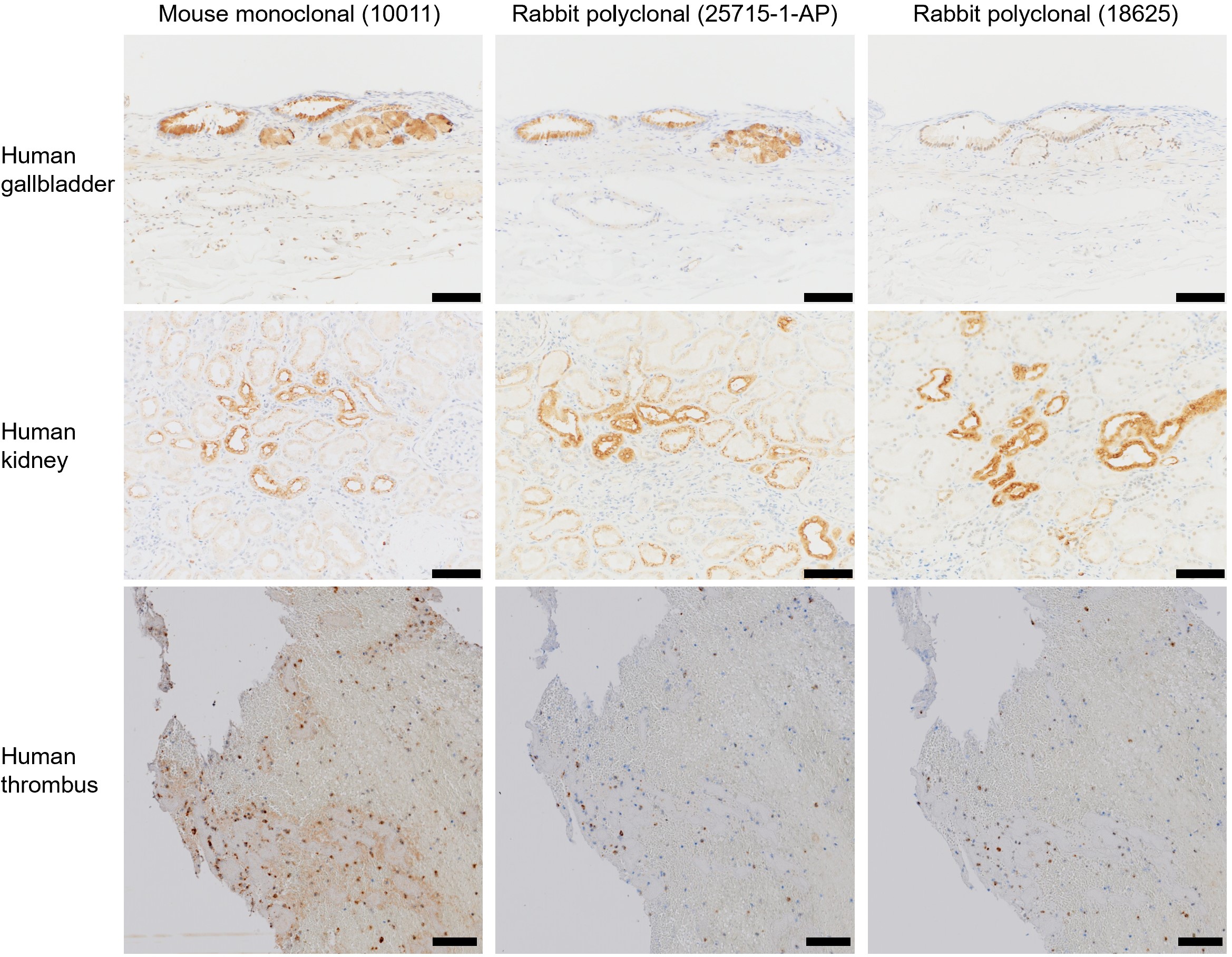

**Fig. S3. Immunohistochemical staining of the positive control samples and thrombi.** Bar = 100 µm.

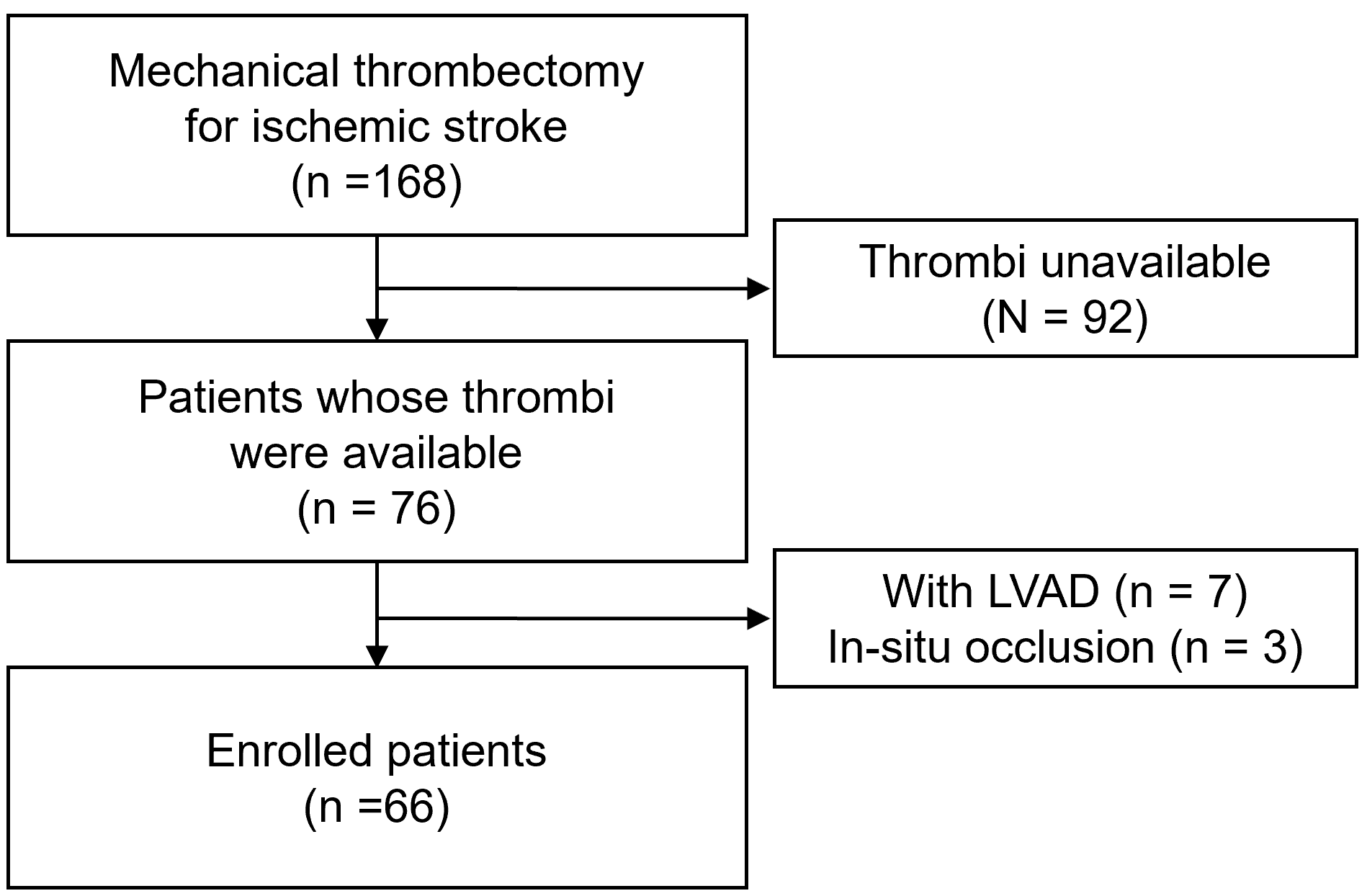

**Fig. S4. Flowchart of the patient selection process.** LVAD, left ventricular assist device.

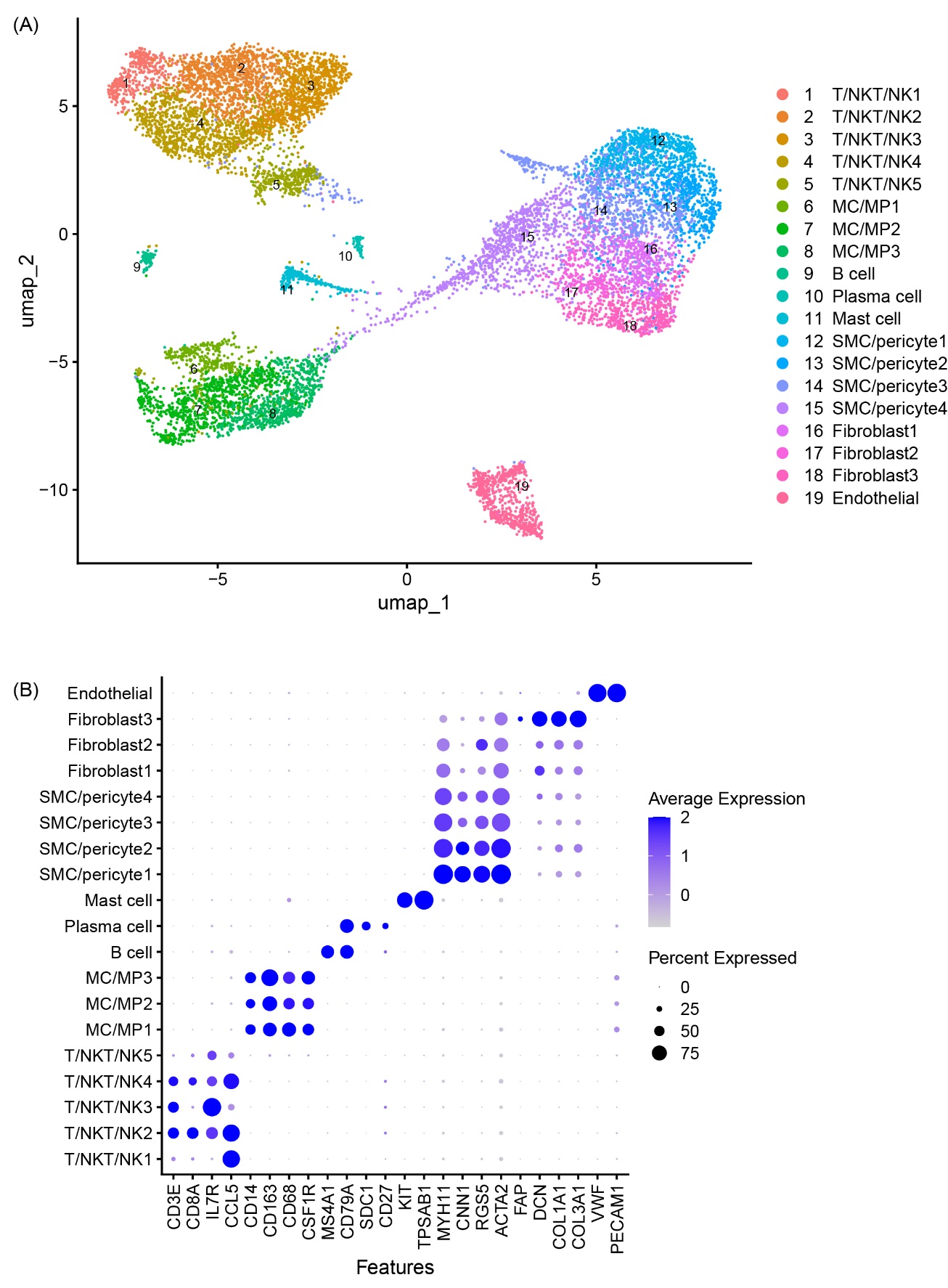

**Fig. S5. Annotation detail. (A)** UMAP plot of the cells recovered from the CTEPH thrombi. Some clusters were manually combined and annotated. **(B)** Signature gene expression as indicated by a dot plot. MC/MP, monocytes/macrophages; T/NKT/NK, T/Natural Killer-like T/Natural Killer; SMC, smooth muscle cells.

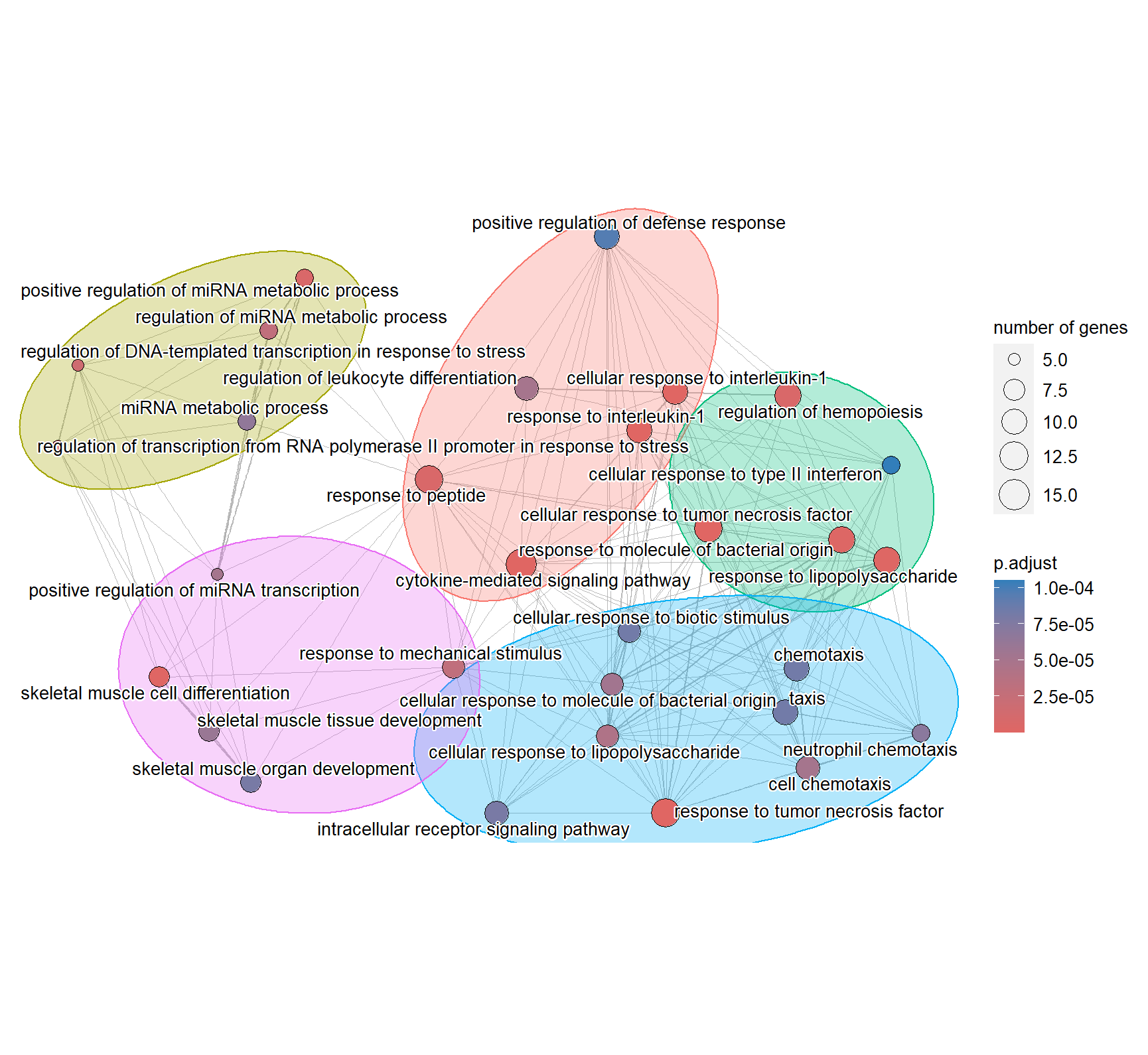

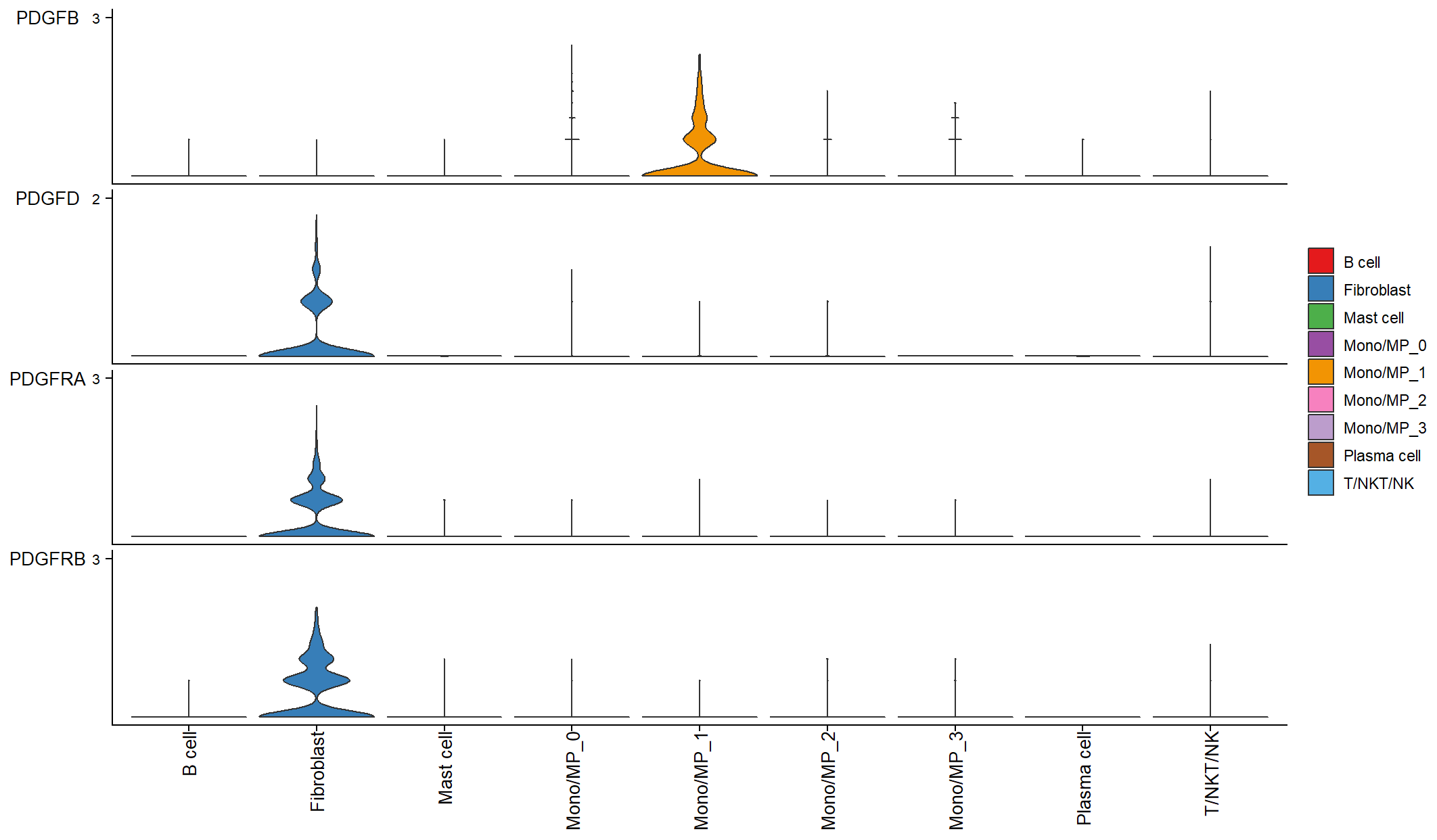

**Fig. S6. Subcluster 1 MC/MPs may contribute to ECM formation in thrombi. (A)** Gene set enrichment analysis results. The upregulated pathways in subcluster 1 MCs/MPs are visualized. **(B)** Expression of the genes in the platelet-derived growth factor subunit β signaling pathway.

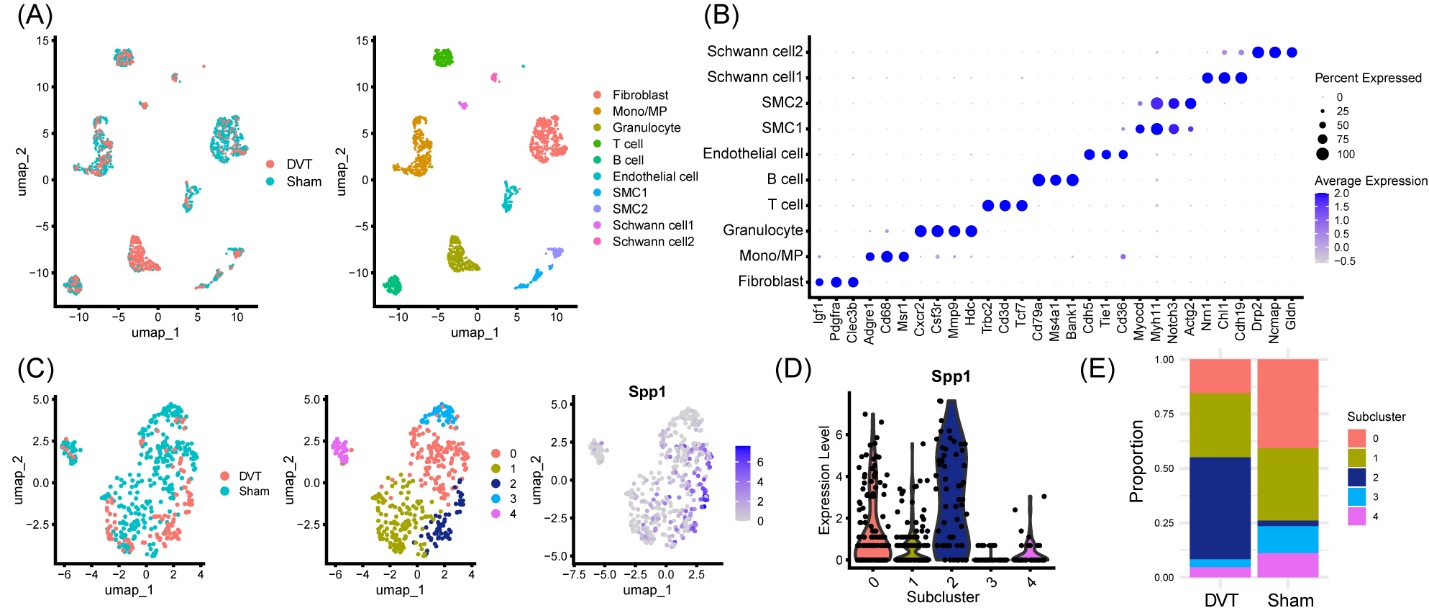

**Fig. S7. The *SPP1*-high monocyte/macrophage subcluster is expanded in the venous wall of mice after thrombosis induction. (A)** UMAP plots of 2,063 cells recovered from the venous vessel wall of mice after deep venous thrombosis (DVT) or sham surgery. **(B)** Expression of marker genes. SMC, smooth muscle cell. **(C)** UMAP plots of 425 monocytes/macrophages classified into five subclusters. **(D)** Violin plot of the *SPP1* expression in each monocyte/macrophage subcluster. Subcluster 2 showed the highest expression of *SPP1* (p_val_adj < 0.001). **(E)** Proportion of each subcluster of monocytes/macrophages according to the indicated conditions. Subcluster 2 (*SPP1*-high) was expanded in the DVT group.

| **Table S1. Background of the subjects** | | | | | | | | | | | | | | |
| --- | --- | --- | --- | --- | --- | --- | --- | --- | --- | --- | --- | --- | --- | --- |
| Number | Age | Sex | Stroke Subtype | mRS | Occluded vessel | NIHSS | Hypertension | Dyslipidemia | Diabetes | Smoking | rt-PA | RIN-blood | RIN-thrombus | Application |
| 1 | 57 | M | CE | 0 | ICA | 18 | Y | Y | N | N | N | 7.2 | 8.2 | RNAseq/qPCR |
| 2 | 90 | F | CE | 4 | MCA | 29 | N | Y | Y | N | N | 7.7 | 6.8 | RNAseq/qPCR |
| 3 | 86 | M | CE | 0 | MCA | 14 | N | N | N | N | N | 7.1 | 4.5 | RNAseq/qPCR |
| 4 | 78 | M | CE | 0 | MCA | 23 | Y | Y | Y | N | N | 7.9 | 2.9 | qPCR |
| 5 | 74 | M | CE | 0 | ICA | 20 | Y | N | N | N | N | 8 | 2.8 | qPCR |
| Abbreviations: CE, cardioembolic; ICA, internal carotid artery; MCA, middle cerebral artery; mRS, modified Rankin Scale; NIHSS, National Institute of Health Stroke Scale; RIN, RNA integrity number; rt-PA, recombinant tissue-plasminogen activator | | | | | | | | | | | | | | |

| **Table S2. Primers used for quantitative real-time PCR analysis.** | | |
| --- | --- | --- |
| **Genes** | **Forward** | **Reverse** |
| *SPP1*_1 | CCGTGGGAAGGACAGTTATG | TAATCTGGACTGCTTGTGGC |
| *SPP1*_2 | GAAGTTTCGCAGACCTGACAT | GTATGCACCATTCAACTCCTCG |
| *CD44* | GATGTCACAGGTGGAAGAAG | CCTTCGTGTGTGGGTAATG |
| *TIMP1* | GACGGCCTTCTGCAATTCC | GTATAAGGTGGTCTGGTTGACTTCTG |
| *TIMP2* | GCACATCACCCTCTGTGACTT | AGCGCGTGATCTTGCACT |
| *ACTB* | CATCCTCACCCTGAAGTACCC | AGCCTGGATAGCAACGTACATG |
